# Epstein-Barr virus miR-BARTs 7 and 9 modulate viral cycle, cell proliferation and proteomic profiles in Burkitt lymphoma

**DOI:** 10.1101/2023.05.19.538556

**Authors:** Brunno Felipe Ramos Caetano, Viviana Loureiro Rocha, Bruno Cesar Rossini, Lucilene Delazari dos Santos, Deilson Elgui de Oliveira

## Abstract

The Epstein-Barr Virus (EBV) encodes viral microRNAs (miRs) that contribute to the pathogenesis of nasopharyngeal and gastric carcinomas, but their potential roles in lymphomas are still to be elucidated. This study sought to assess the impact of knocking down EBV miRs BART 7 and BART9 in EBV-positive Akata cell lines using CRISPR/Cas9 technology. Compared to cells harboring the wild-type (WT) EBV genomes, Akata cells subjected to CRISPR/Cas9-mediated knockdown of EBV BART 7 and BART9 showed a significant reduction in the expression of viral miRs, confirming the validity of the experimental model. Knocking down both BART7 and BART9 caused a significant reduction in cell viability and proliferation rates while increasing the expression of EBV lytic genes. Global proteomic analysis shows that knocking down EBV BART7 significantly decreased the expression of ubiquitin/proteasome proteins while increasing RNA binding proteins (RBPs). On the other hand, BART9 knockdown caused a decrease in proteins associated with oxidoreductase activity, including the metabolism of fatty acids. Our results unravel previously unknown roles for EBV miRs BART7 and BART9 on cellular pathways relevant to both viral biology and lymphomagenesis.

## 1. Introduction

The Epstein-Barr Virus (EBV) is a ubiquitous gammaherpesvirus implicated in the pathogenesis of lymphoproliferative disorders and several human cancers. These include the undifferentiated form of nasopharyngeal carcinoma (NPC), Burkitt lymphoma (BL), classical Hodgkin lymphoma (cHL), various non-Hodgkin lymphomas (NHL), and a small fraction of gastric carcinomas[1]. Epithelial and B-cells are the primary targets of EBV infection, and the virus latently infects memory B cells to establish a life-long, persistent infection in human hosts. Several viral products expressed during the EBV latent cycle have known oncogenic activities and contribute to EBV-associated malignancies, including the latent membrane proteins (LMPs) 1 and 2, the Epstein–Barr virus nuclear antigen 1 (EBNA1), the Epstein-Barr virus-encoded small RNAs (EBERs), and, more recently, EBV-encoded microRNAs (miRs)[2].

EBV was the first human virus identified to encode miRNAs, clustered in two regions of the viral genome: the BamHI fragment H rightward open reading frame 1 (miRs-BHRF) and the BamHI-A rightward transcripts (miRs-BART)[3]. EBV encodes 25 precursor miRs (pre-miRs), producing 48 mature viral miRs[4]. The EBV miRs-BART locate within intronic segments of the genomic locus encoding the BART transcripts, an extensively-spliced region expressed in all forms of viral latency programs. Although EBV-encoded miRNAs are not required to produce new virions, they play critical roles in regulating the viral lytic cycle and the pathogenesis of some EBV-associated cancers[5,6].

The EBV miRs-BART are highly expressed in nasopharyngeal and gastric carcinomas, contributing to malignant transformation and tumor progression[7]. Conversely, miR-BART expression levels in lymphoid malignancies are lower than that found in epithelial cells., and their impact on lymphomagenesis is not sufficiently elucidated[8]. In 2011, Ramakrishnan and colleagues reported that the expression of EBV miR-BART9 was associated with increased levels of LMP1 (both mRNA and protein), promoting cell proliferation in nasal NK/T-cell lymphomas (NKTL)[9]. Furthermore, EBV miRs-BART 7 and 9 reportedly reduce the expression of the *BCL6* gene in EBV-associated Diffuse Large B cell Lymphoma (DLBCL), implicated in the modulation of cell cycle and differentiation, apoptosis, and the levels of local inflammation[10]. Therefore, in this study, we sought to assess the expression of EBV miRs BART7 and BART9 in a panel of lymphoid cells constitutively infected by EBV and investigate the in vitro effects of knocking down these viral miRNAs in EBV-positive Akata cells.

## 2. Material and Methods

### 2.1 Cell lines and other resources

Akata, MutuI, and P3HR1 cells were generously provided by Professor Benjamin Gewurz (Harvard Medical School, Harvard U., MA, USA), while BC1, BC2, BC3, Daudi, IBL-1, Jiyoye, and Rael were a gift from Professor Ethel Cesarman (Weill Medical College, Cornell U., NY, USA). All cell lines were cultured in RPMI 1640 (Thermo Fisher Scientific, Waltham, MA, USA) supplemented with 10% Fetal Bovine Serum (FBS; Sigma-Aldrich, St Louis, MO, USA) and 0.4% gentamicin (Thermo Fisher), incubated at 37°C in 5% CO_2_. EBV-positive Akata cell lines with stable Cas9 (*Streptococcus pyogenes*) expression were generated by lentiviral transduction and blasticidin selection. For the stable selection of EBV-positive Cas9-expressing clones, Akata cells were cultured in a medium supplemented with 500 μg/mL geneticin (Thermo Fisher) and 10 μg/mL blasticidin (Thermo Fisher). All cells were confirmed to be free of mycoplasma (PCR-based method) and authenticated by Short Tandem Repeat (STR) analysis using the GenePrint® 10 System (Promega, Madison, WI, USA; Supplementary Material, Table S1). A list of other experimental resources and reagents used in this study can be found in Supplementary Material Table S2.

### 2.2 Design and synthesis of CRISPR/Cas9 single-guide RNAs (sgRNAs)

The sgRNAs sequences to target EBV miRs-BART 7 and 9 were selected using the CRISPick online tool (Broad Institute, https://portals.broadinstitute.org/gppx/crispick/public) using hairpin sequences retrieved from miRbase (http://www.mirbase.org). Customized DNA oligonucleotides with additional nucleotides inserted for restriction enzyme-mediated cloning (sequences in Supplementary Material, Table S3) were synthesized by Thermo Fisher. T4 ligase-mediated annealing of sgRNAs resuspended in Tris-EDTA buffer (TE, 10 mM Tris, 0.1 mM EDTA, pH 8.0) was performed using the T4 Ligase Reaction Buffer (New England Biolabs, NEB, Ipswich, MA, USA).

### 2.3 Production of lentiviral vectors for sgRNA expression

For the CRISPR/Cas9-mediated genomic edition of EBV miRs-BART 7 and 9, the recombinant vectors LentiGuide-Puro-BART7 and LentiGuide-Puro-BART9 were constructed by cloning the respective sgRNAs into the LentiGuide-Puro backbone (RRID: Addgene_52963, http://n2t.net/addgene:52963), a gift from Feng Zhang[11]. Briefly, the backbone vector was digested with BsmBI (New England Biolabs), followed by gel purification using the QIAquick PCR & Gel Cleanup Kit (Qiagen, Valencia, CA, USA). The linearized vector and the sgRNAs heteroduplexes were ligated overnight at 16°C using T4 DNA ligase (NEB), and the ligation product was used to transform One Shot™ Stbl3™ cells (Thermo Fisher). Transformed colonies were subjected to plasmidial isolation, and the constructs were verified by Sanger sequencing using the hU6.F primer.

Lentiviral particles harboring the sgRNAs for CRISPR/Cas9 edition of EBV miRs-BART 7 and 9 were obtained after transfection of HEK293FT cells with the LentiviralPuro-based constructs and the packaging vectors psPAX2 (RRID: Addgene12260; http://n2t.net/addgene:12260) and VSVG (RRID: Addgene_14888; http://n2t.net/addgene:14888). The transfection was performed with Lipofectamine™ 3000 (Thermo Fisher) in 6-well plates. DNA-liposome complexes were prepared in OptiMEM™ (Gibco), homogenized, and added dropwise into each well containing attached HEK293FT cells cultivated in DMEM medium (Gibco). After incubation for 24h at 37°C in 5% CO_2_, the transfection media was replaced with fresh RPMI. The lentiviral particles were collected at 48 and 72 h post-transfection, filtered through 0.45 μM PES syringe filters, and directly used for lentiviral transduction.

### 2.4 Lentiviral transduction in EBV-infected Akata cells expressing Cas9

EBV-positive Akata cells expressing Cas9 (Akata-EBV/Cas9) were transduced with fresh stocks of lentiviral particles harboring the sgRNA vectors for the CRISPR-Cas9 edition of EBV miRs BART 7 and 9. After 48h post-transduction, the cells received fresh RPMI medium supplemented with 10 μg/mL puromycin (Invitrogen) for eukaryotic selection. Cells stably transduced obtained after 2 weeks were then propagated and maintained in culture with RPMI medium supplemented with 10% FBS, 1 μg/mL puromycin, 500 μg/mL geneticin, and 10 μg/mL blasticidin.

### 2.5 T7 endonuclease I assay and TIDE analyses

The T7 endonuclease I (T7EI) assay was used to assess the efficiency of the CRISPR/Cas9 edition of the EBV genome[12]. Specific primers for amplification of a region flanking the predicted target sites for CRISPR/Cas9 edition were designed and used for PCR amplification (Supplementary Material, Table S3) with the Platinum™ High Fidelity Taq DNA Polymerase (Thermo Fisher). The PCR product was separated using 1.5% agarose gels and purified using the QIAquick PCR & Gel Cleanup Kit (Qiagen). Purified PCR products (300 ng) were then denatured at 95°C for 5 min and cooled at room temperature to form heteroduplexes using NEBuffer2.1 1x (NEB). The T7 endonuclease I (NEB) was then added to the heteroduplexes and incubated for 30 min at 37°C. DNA fragments were separated on 1% agarose gel, and the cleavage products were evaluated using a Reverse Mass DNA Ladder in Gel Loading Dye Orange (NEB).

Following the previously published protocol[13], we used the Tracking of Indels by DEcomposition (TIDE) analysis to determine the spectrum and frequency of targeted mutations generated in Akata-EBV/Cas9 cells by CRISPR/Cas9. Genomic DNA was amplified, the PCR product was purified as described previously, then subjected to Sanger sequencing using the 3500 Series Genetic Analyzer (Thermo Fisher) device. The samples sequenced in parallel using the forward and reverse primers (<u>Table S2</u>) had their sequence trace files uploaded to the online TIDE tool (https://tide.nki.nl) for analysis of the indel/HDR spectrum.

### 2.6 Assessment of expression of human and viral genes by qPCR

Total RNA was isolated using the TRIzol™ (Invitrogen) reagent, followed by purification with the Qiagen RNeasy column (Qiagen), treatment with DNAse (Sigma-Aldrich), and quantification by spectrophotometry using the NanoDrop® device (Thermo Fisher). The complementary DNA (cDNA) was synthesized using the High-Capacity cDNA Reverse Transcription Kit (Thermo Fisher), according to the manufacturer’s instructions. Custom reverse transcription primers (Canopy Biosciences, St. Louis, MO, USA) were added to total RNA samples before cDNA synthesis to amplify the EBV miRs-BART under evaluation.

The assessment of gene expression was performed by quantitative real-time Polymerase Chain Reaction (qPCR) using the GoTaq® qPCR Mix (Promega) in the 7500 Fast Real Timer PCR system (Applied Biosystems®). SNORD47 and SNORD48 or HSP90 and RPS13 were used as reference endogenous controls to assess the expression of miRs and mRNAs, respectively. The relative expression levels were estimated using the comparative Ct method[14]. The expression of the EBV lytic proteins BZLF1 (i.e., Zta, NCBI gene ID: 3783744) and BLLF1 (i.e., gp350, NCBI gene ID: 3783713) were assessed in parental Akata-EBV/Cas9 (wt) and its EBV miRs-BART mutants. The primers sequences and qPCR conditions for these assays are indicated in the Supplementary Material (Table S3).

### 2.7 Luciferase reporter assays

The activity of EBV miRs-BART 7-3p and 9-3p was evaluated in Akata-EBV/Cas9 cells by luciferase reporter gene assay with constructs based on the psiCheck-2 backbone (Promega). 3’-UTR sequences with miRNA-binding sites (Supplementary Material, Table S3) and the respective negative controls (scramble, SCR) were cloned into the psiCheck-2 vector according to the manufacturer’s instructions. Furthermore, a luciferase reporter assay to assess the NFκB activity was performed using a pcDNA3.1-NFκB-luc (firefly luc; a gift of Prof. Ethel Cesarman), and the pGL4.73 vector (hRluc/SV40, renilla; Promega). The pcDNA3.1-NFκB-luc contains a sequence reporter with the luciferase gene under the control of a kappa enhancer element. Akata-EBV/Cas9 cells expressing psiCheck-2-BART7-3p, psiCheck-2-BART9-3p, pcDNA3.1-NFκB-luc, and the pGL4.73 were generated by transfection with Lipofectamine™ 3000 (Thermo Fisher). The firefly and renilla luciferase activity assessment were conducted 48h post-transfection using the Dual-Glo® Luciferase Assay System (Promega), according to the manufacturer’s protocol and the GloMax® Discover Microplate Reader (Promega).

### 2.8 Cell viability and proliferation assays

An MTS assay using the CellTiter 96® Aqueous One Solution (Promega, Madison) was performed to assess the rates of cell viability and proliferation in vitro. Briefly, 2×10 ^3^ cells were seeded onto 96-well plates in complete medium and 20 μL of the CellTiter reagent. After incubation, the formazan dye produced by viable cells was quantified by measuring the absorbance at 490 nm on the Bio-Rad Model 680 microplate reader (Bio-Rad Laboratories, Hercules, CA, USA).

### 2.9 Proteomic analysis

Whole-cell lysates from Akata-EBV/Cas9 (wt) and miRs-BART 7 and 9 mutants were extracted using a detergent-free buffer containing 8 M urea, 75 mM NaCl, 50 mM Tris-HCl pH 8.2, 50 U/mL benzonase, 2 mM MgCl_2,_ and protease inhibitors (cOmplete™, EDTA-free Protease Inhibitor Cocktail; Roche Life Science, Indianapolis, IN, USA), as previously reported[15]. Briefly, cell pellets were washed twice with DPBS, resuspended in lysis buffer (100 μL in 1×10 ^6^ cells), and sonicated (30 sec, 5 sec on, and 5 sec off) at 20% amplitude with the Vibra-Cell™ VCX 750 (Sonics & Materials Inc., Newtown, CT, USA). Samples were then incubated at 4°C for 30 min and centrifuged at 16,000 x g for 30 min at 4°C. The supernatant was collected and quantified using the Bio-Rad Protein Assay Kit and the Quick Start Bovine Serum Albumin Standard Set (Bio-Rad) and stored at −80°C before use.

Protein samples (50 μg) were diluted in 50 mM ammonium bicarbonate buffer (AmBic, NH_4_HCO_3_; Sigma-Aldrich), 0.2% (m/v) RapiGest SF (Waters™, Milford, MA, USA), and incubated for 60 min at 37°C. After surfactant incubation, 100 mM dithiothreitol (DTT, 2.5 μL; Sigma-Aldrich) and 300 mM 2-iodoacetamide (IAA, 2.5 μL; Sigma-Aldrich) were added to the samples, followed by incubation for 30 min at room temperature. Samples were then digested with Sequencing Grade Modified Trypsin (0.1 μg/μL, Promega) for 16 h at 37°C. Upon digestion, reactions were acidified with 5% trifluoroacetic acid (TFA, v/v) and incubated for 90 min at 37°C to stop proteolysis. Samples were centrifuged at 16,000 x g at 4°C for 30 min to remove insoluble debris. The supernatant containing soluble peptides was collected and dried overnight using the Vacufuge® Concentrator Plus (Eppendorf, Hamburg, HH, Germany). Peptides were desalted, purified, and concentrated using the Pierce™ C18 Spin Columns (Thermo Fisher) according to the manufacturer’s instructions.

### 2.10 Liquid chromatography-tandem mass spectrometry (LC-MS/MS)

Mass spectrometry was performed to identify and quantify differentially expressed proteins in Akata-EBV/Cas9 miRs-BART 7 and 9 mutants. The obtained peptides were subjected to MS, followed by sequencing analysis using a liquid chromatography-tandem mass spectrometry system consisting of the UltiMate 3000 LC liquid nanochromatography system (LC Packings DIONEX, Sunnyvale, CA, USA) coupled to the Q-Exactive™ (Thermo Fisher). LC-MS/MS was conducted according to the parameters previously described[16]. Expressed proteins were identified with the PatternLab software version 4.0.0.84 using the UniProt proteome of *Homo sapiens* (UniProt ID: UP000005640) as reference^17^. Fragment mass tolerance was set to 40 ppm with a false discovery rate (FDR) of ≤1%. Identified proteins were submitted to parsimony analysis, and quantitation analysis using extracted-ion chromatogram (XIC) ^17^ was processed using the MetaboAnalyst (https://www.metaboanalyst.ca) web-based tool, with FDR of ≤5%. Also, we used the RawVegetable software for quality control of raw MS files[18,19].

## 3. Results

### 3.1 CRISPR/Cas9-targeted mutagenesis efficiently knockdowns the expression of EBV miRs-BART

The BART7 and BART9 are located within the intronic regions of the BART cluster 2 in the viral genome (**Fig. 1a**). Two synthetic guide RNAs (sgRNAs) for each miR-BART were designed to target different regions within the 5’-Dicer processing site of the precursor viral miRNA (pre-miRNA) sequences to disrupt the generation and maturation of the targeted EBV miR-BART using CRISPR/Cas9 technology (**Fig. 1b** and **1c**). Akata-EBV/Cas9 cells transduced with the lentiviral vectors (LentiGuide-Puro-BART7 and LentiGuide-Puro-BART9) were evaluated for insertion and deletion mutations (*indels*) caused by CRISPR/Cas9 edition of the EBV genome.

**Figure 1.**
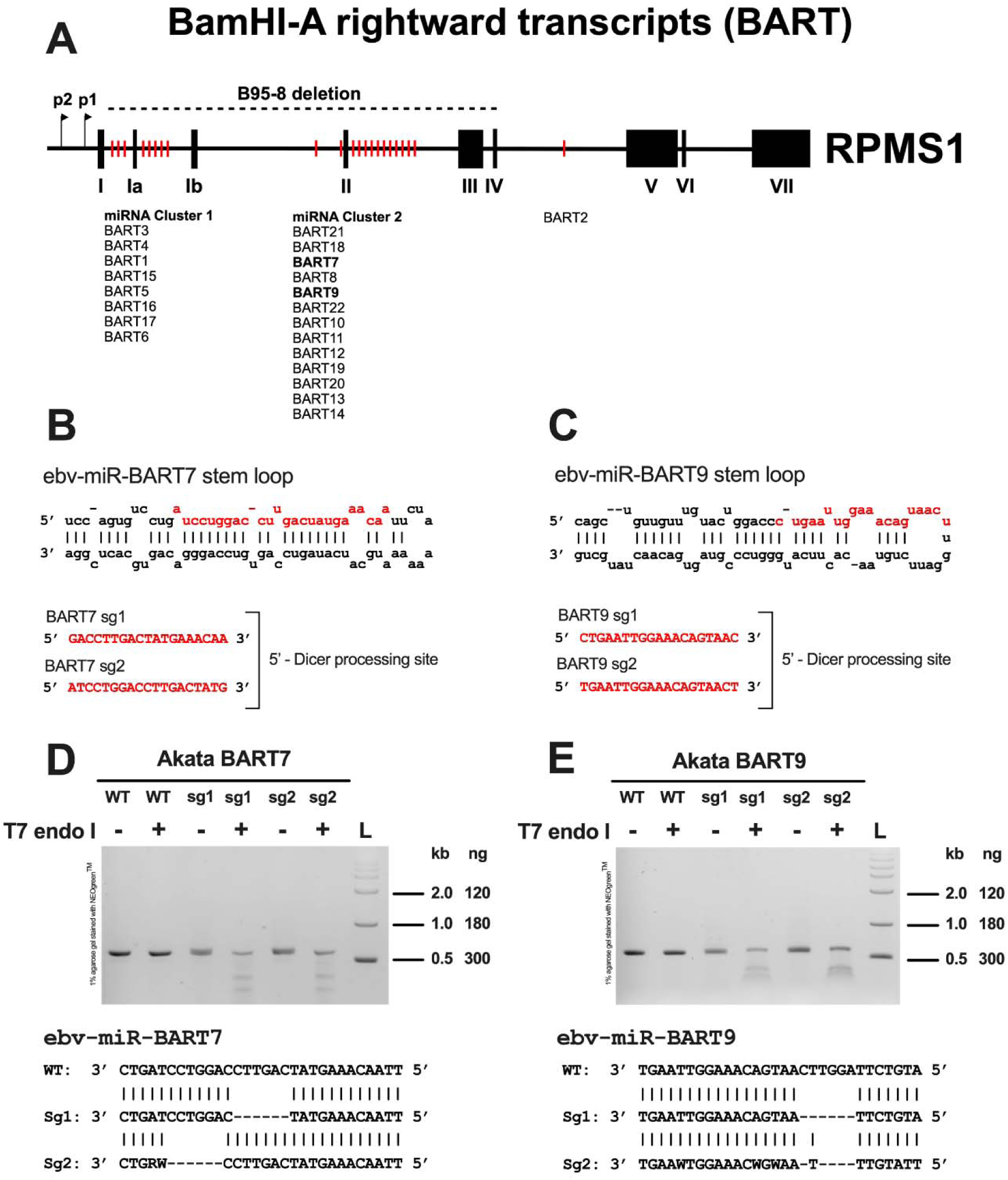
(a) Schematic representation of the EBV miRs-BART clusters within the intronic region of the BART transcripts. Sequences of the pre-miRNAs and sgRNAs designed for Cas9/sgRNA-directed mutagenesis of EBV miR-BART7 **(b)** and miR-BART9 **(c)**. Both sgRNAs were designed to target the 5’-Dicer processing site, impairing the maturation of pre-miRNAs. **(c** and **d)** T7EI assay and sequence analysis of Akata WT and mutants after transduction with sgRNAs and stable selection.

The T7 endonuclease I mismatch detection assay revealed two expected cleavage bands (300 and 200 bp) for both sgRNAs, identifying the indels generated in the EBV genomes (**Fig. 1d** and **1e**). DNA sequencing of a flanking region encoding ebv-miR-BART7 and ebv-miR-BART9 also showed several deletions within the predicted sites (**Fig. 1d** and **1e**). The *indels* profile obtained by TIDE analysis showed that the most frequent mutations found in the viral DNA of Akata-EBV/Cas9 cells were deletions spanning 1-6 nucleotides, indicating a CRISPR/Cas9 genome edition efficiency above 70% for EBV miRs-BARTs (Supplementary Material, **Fig. S1**).

To better understand how CRISPR/Cas9 edition affected the expression of the investigated EBV miRs-BART, we assessed the expression BART7-3p and BART9-3p in both Akata-EBV/Cas9 wild type (WT) and mutants (ΔBART7 and ΔBART9) using RT-qPCR. As shown in **Fig. 2a,** a significant decrease in the expression of ebv-miR-BART7-3p and ebv-miR-BART9-3p was found in their respective mutants, with higher knocking down levels observed for Akata-EBV/Cas9 with sg2-mediated EBV miRs-BART edition (ΔBART7sg2 and ΔBART9sg2). The knockdown of BART7 and BART9 in the Akata-EBV/Cas9 cells was further verified using luciferase assays with cells expressing the reporter constructs psiCHECK-2-BART7-3p and psiCHECK-2-BART9-3p, harboring the 3’-UTR sequences targeted by the respective viral miRNAs (**Fig. 2b**). As shown in **Fig 2c,** Akata-EBV/Cas9 cells harboring EBV mutants ΔBART7 and ΔBART9 showed increased luciferase activity when compared to the EBV-WT control, and the highest levels of abrogation of the EBV miRs-BART 7 and 9 were observed for cells infected with EBV mutants ΔBART7sg2 and ΔBART9sg2, respectively.

**Figure 2.**
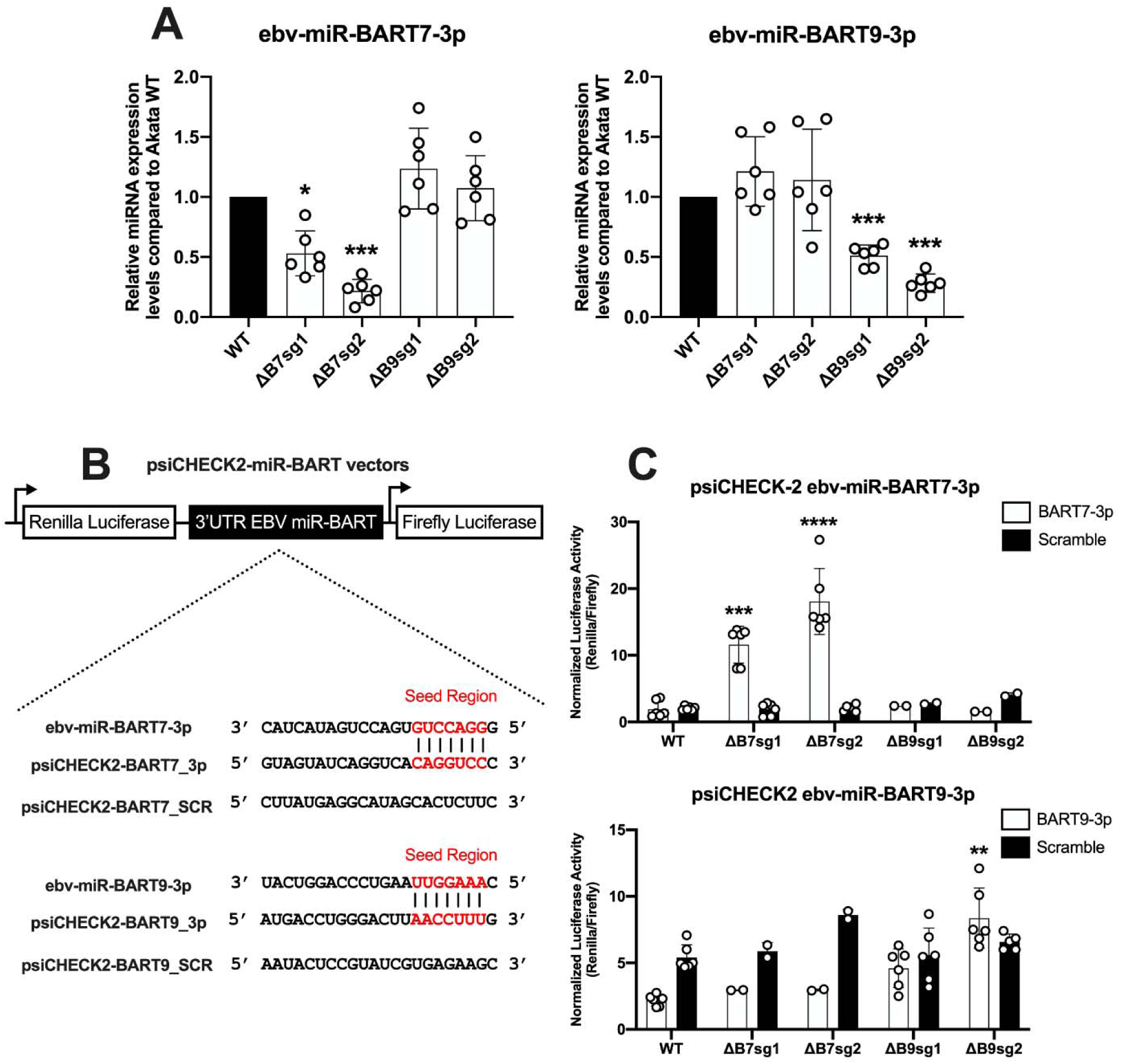
Effects of CRISPR/Cas9-targeted mutagenesis on the expression of EBV miR-BART7 and miR-BART9 in Akata mutants. **(a)** Relative expression of EBV miR-BART7-3p and miR-BART9-3p in Akata mutants assessed by RT-qPCR. The human miRNAs Snord47 and Snord48 were used as endogenous controls. *p=0.0003 and ***p<0.0001 when compared to Akata WT; unpaired two-sample t-test. **(b)** Mechanism of psiCheck-2 luciferase reporter vector. **(c)** Luciferase activity in Akata WT and mutants transfected with report vectors for EBV miR-BART7-3p and miR-BART9-3p. **p=0.00017 when compared to Akata WT; one-way ANOVA followed by Dunnet’s test.

### 3.2 EBV miRs-BART knockdown reduces cell viability and proliferation

We assessed the effects of knocking down BART7 and BART9 on the cell viability and proliferation rates of Akata-EBV/Cas9 cells infected with EBV ΔBART7 or ΔBART9 compared with cells infected with EBV-WT (control). As shown in **Fig 3a** and **3b**, knocking down both viral miRNAs caused a significant reduction in cell viability and proliferation in cells constitutively infected with the EBV mutants compared to the control.

**Figure 3.**
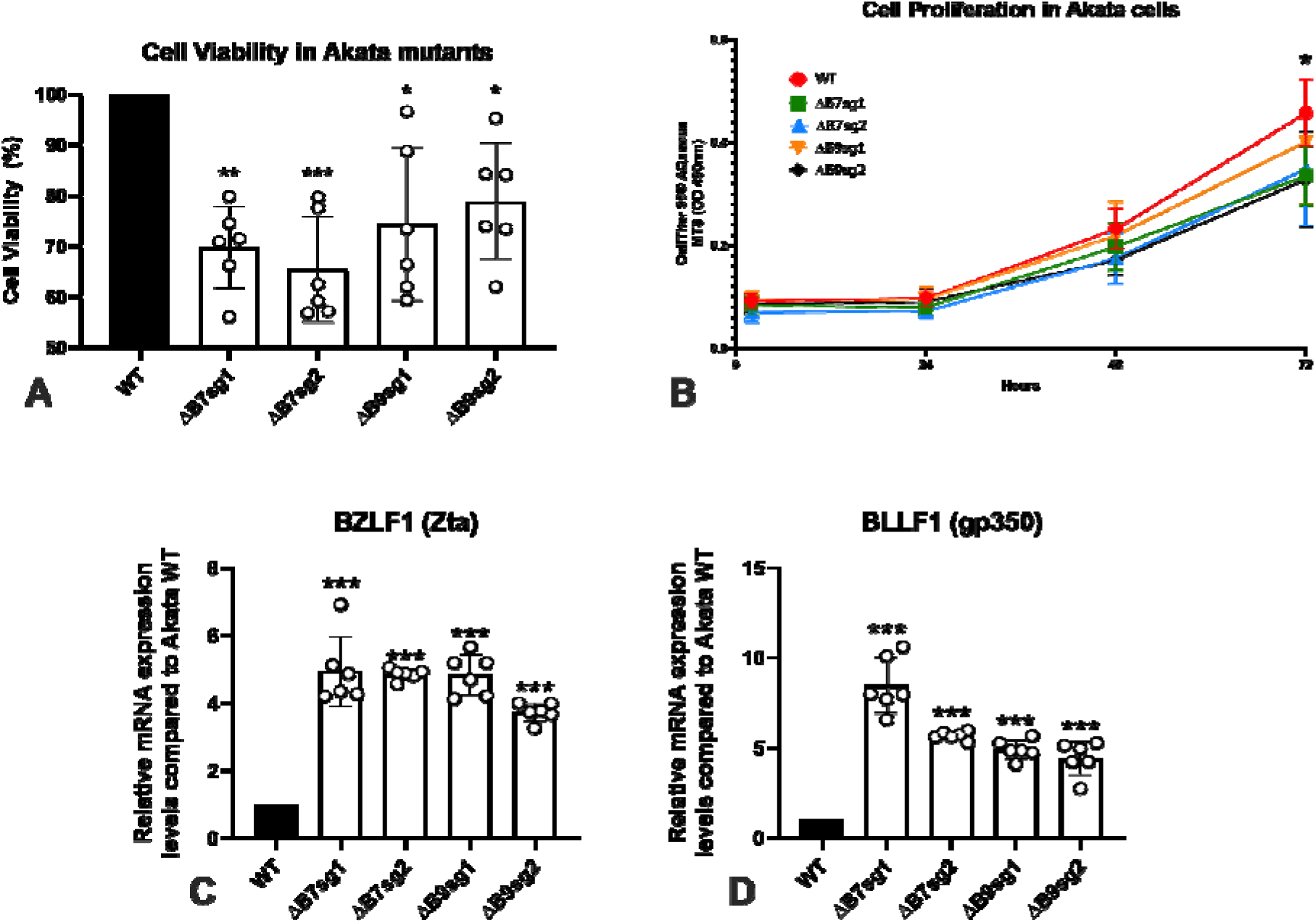
Cell viability and proliferation in Akata WT and mutants assessed by the CellTiter 96 ® AQueous One Solution Cell Proliferation Assay. **(a)** Cell viability in Akata WT and mutants; *p=0.0001, **p=0.0009 ***p<0.0001 when compared to Akata WT cells; one-way ANOVA followed by Dunnet’s test. (**b**) Cell proliferation in Akata WT and mutants following 4, 24, 48, and 72h after plating. *p=0.0320 when compared to Akata WT; one-way ANOVA followed b y Dunnet’s test. Expression of the viral lytic genes **(c)** BZLF1 (Zta) and **(d)** BLLF1 (Rta) in Akata mutants by RT-qPCR. Expression of human immune checkpoint genes **(c)** TIM3 and **(d)** PDLF1 by RT-qPCR. ***p<0.0001 when compared to Akata WT; unpaired two-sample t-test.

### 3.3 Knocking-down EBV miRs-BART induces lytic cycle genes

EBV BART miRNAs modulate critical pathways related to the viral lytic cycle, with consequences for viral persistency and immune escape [20]. Based on this, we assessed the expression levels of the EBV early-immediate lytic gene BZLF1 (Zta, gene ID: 3783744) and the late lytic gene BLLF1 (gp350, gene ID: 3783713) in Akata-EBV/Cas9 cells infected with EBV-WT and mutants by RT-qPCR. Akata cells with EBV mutants ΔBART7 and ΔBART9 showed an increase in the expression of both BZLF1 and BLLF1 lytic genes compared to cells infected with the EBV-WT control (**Figs. 3c** and **3d**).

### 3.4 Identification of proteomic profiles in ebv-miRs-BART mutants

To further characterize the effects of knocking down the EBV BART 7 and BART9 in our experimental model of Akata-EBV/Cas9 cells, we investigated the global expression profile of proteins of the cells using LC-MS/MS, followed by pathway enrichment (PE) analyses based on proteins uniquely-expressed in cells infected with mutants for ebv-miR-BART7 or ebv-miR-BART9 compared to cell line infected with EBV-WT (control).

Overall, 2,118 and 2,065 proteins were identified in ΔBART7sg2 and ΔBART9sg2 mutants, respectively (Supplementary Material, Datasets 1 and 2). Proteins found to be significantly up or down-regulated were 59 in cells infected with EBV ΔBART7sg2 and 44 for cells infected with EBV ΔBART9sg2, when compared to the control (**Tables 1** and **2**). Proteins involved in RNA-binding, nucleotide biosynthesis and nucleotide metabolism pathways were enriched in cells infected with the EBV ΔBART7sg2 mutant, while cells infected with EBV ΔBART9sg2 mutant showed enrichment of proteins associated with oxireductase and catalytic activities (**Fig.4**).

**Figure 4.**
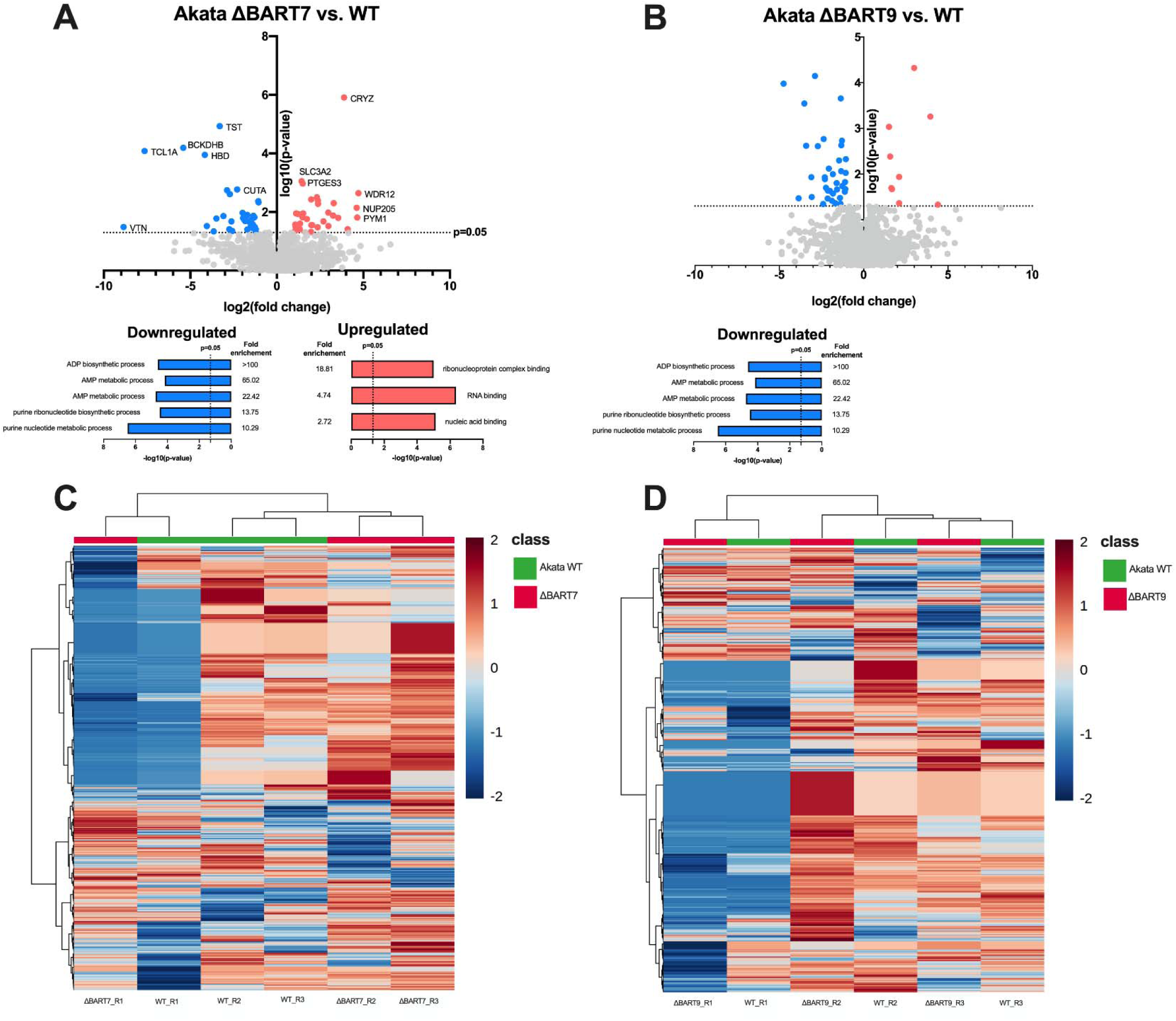
LC/MS results of differently expressed proteins in Akata-EBV/Cas-9 cells with knockdown of ebv-miR-BART7 and ebv-miR-BART9 expression by CRISPR/Cas9 genomic edition, compared to the parental, non-edited cell line. **(a)** and **(b)** Volcano plots and enrichment analysis of significantly up- and down-regulated proteins. **(c)** and **(d)** Clustering results of differentially expressed proteins, shown as heatmap. Distance and clustering measured using ward.D algorithm. Proteins were considered regulated if fold-change values were ≥1.5 or ≤0.5 (log2 FC) with p-value < 0.05 (-log10[p]).

**Table 1.**
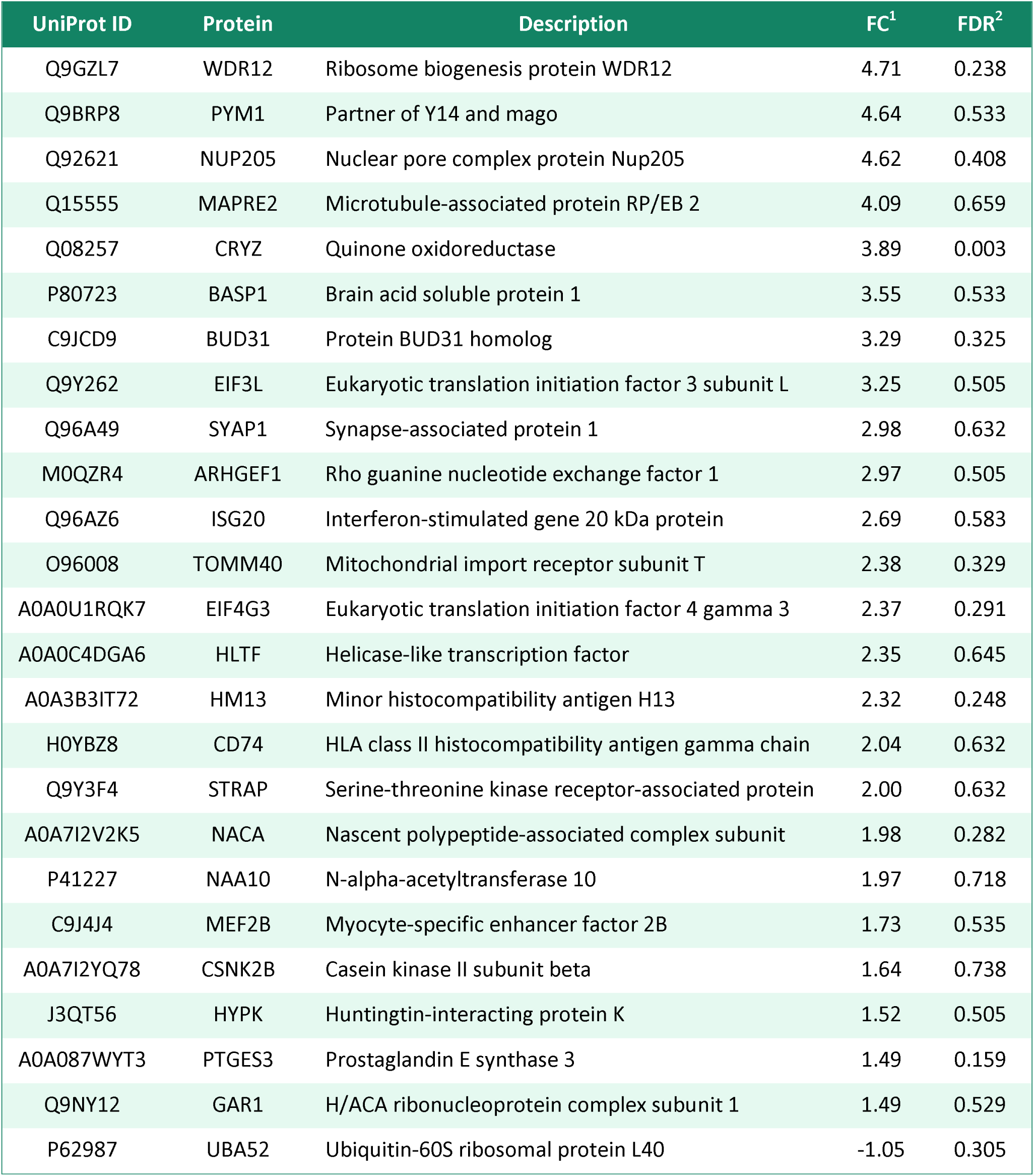
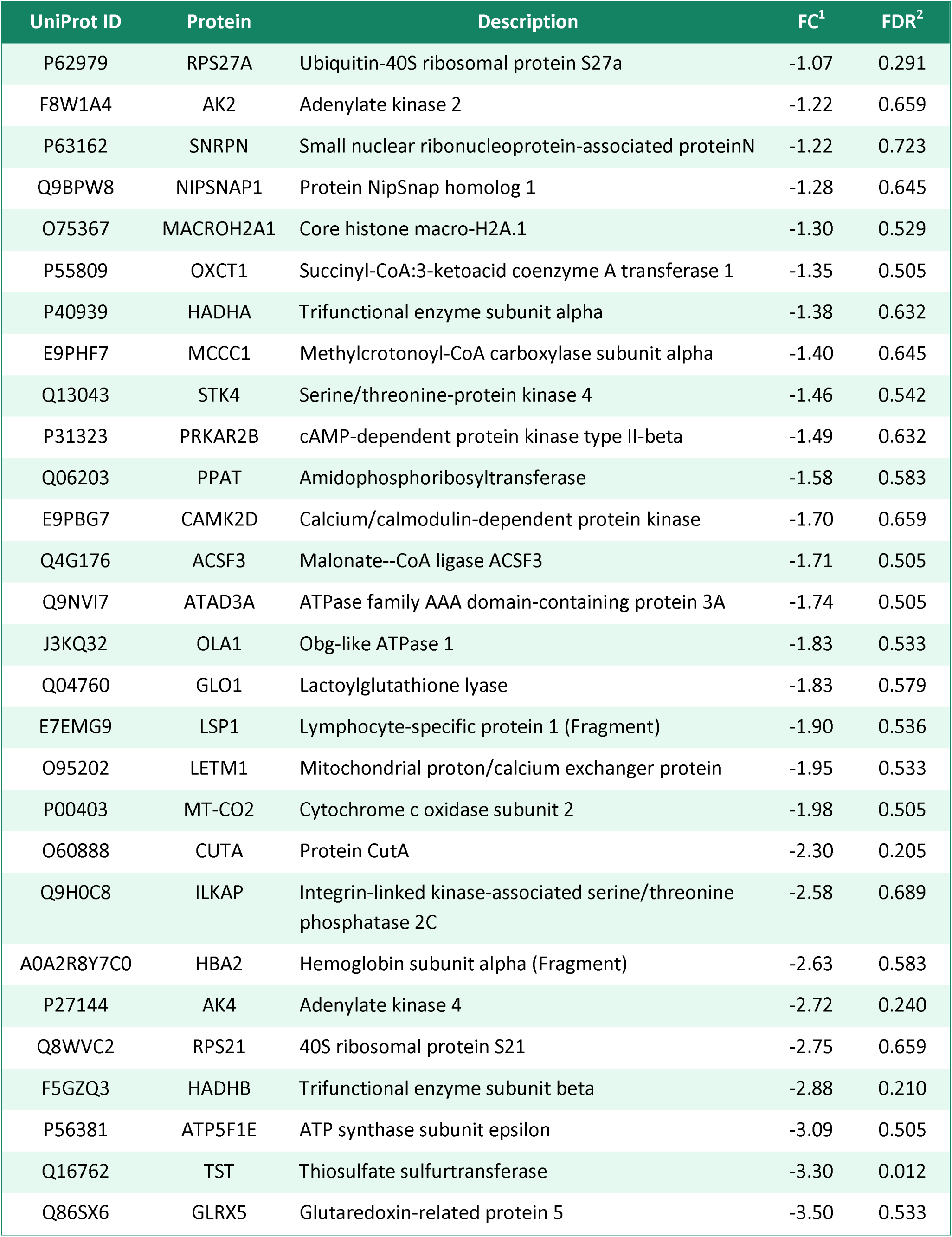
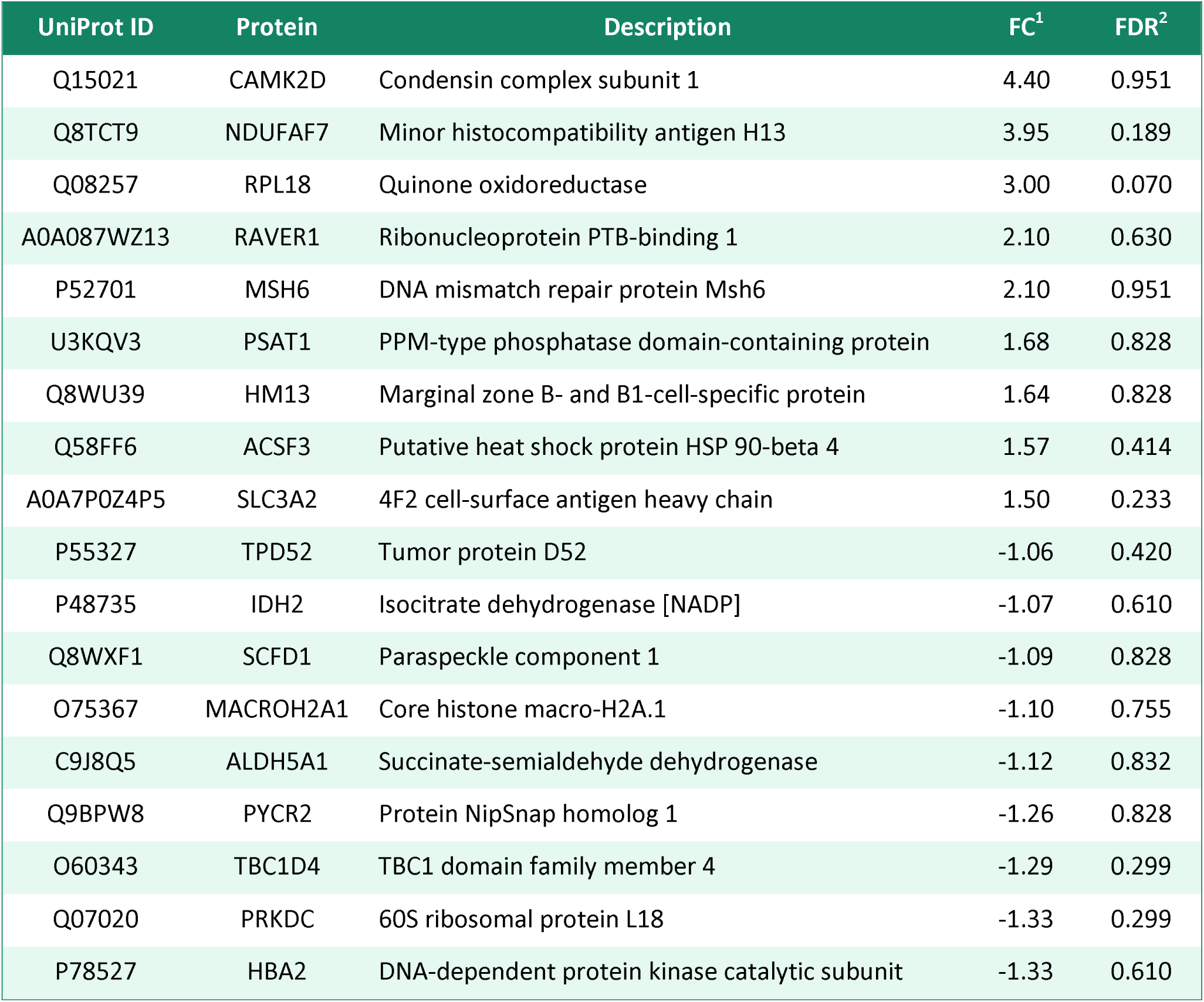
List of proteins significantly up- and down-regulated in ΔBART7_sg2 mutant compared to the Akata-EBV/Cas9 wild-type.

**Table 2.**
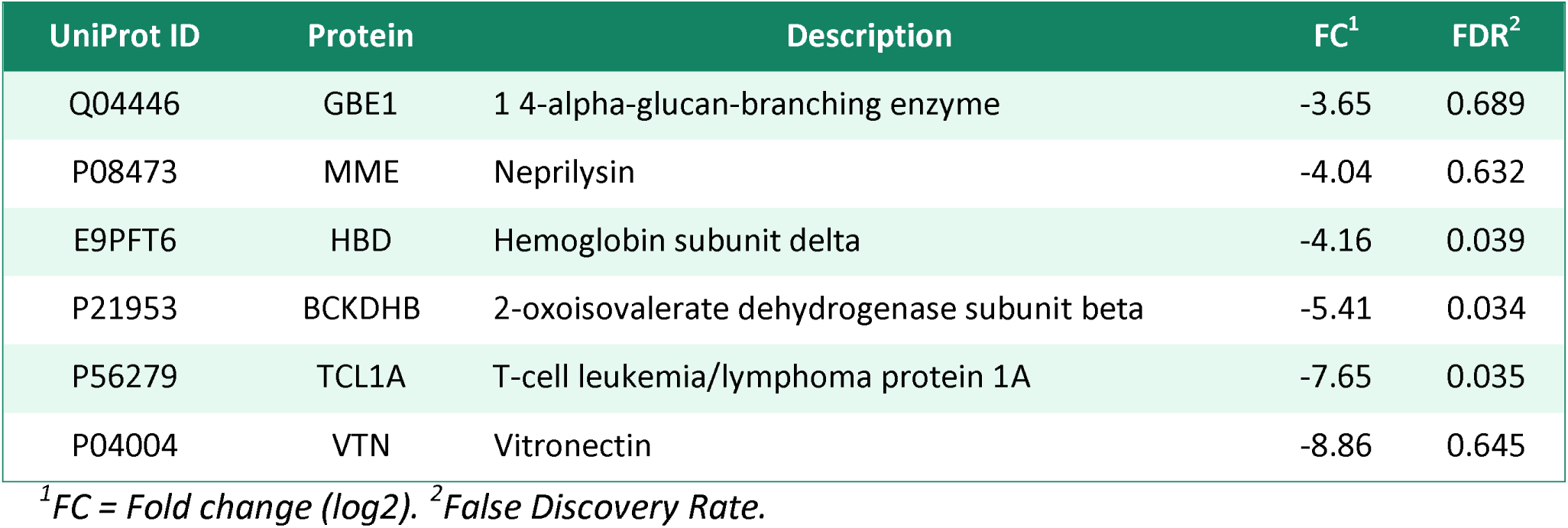
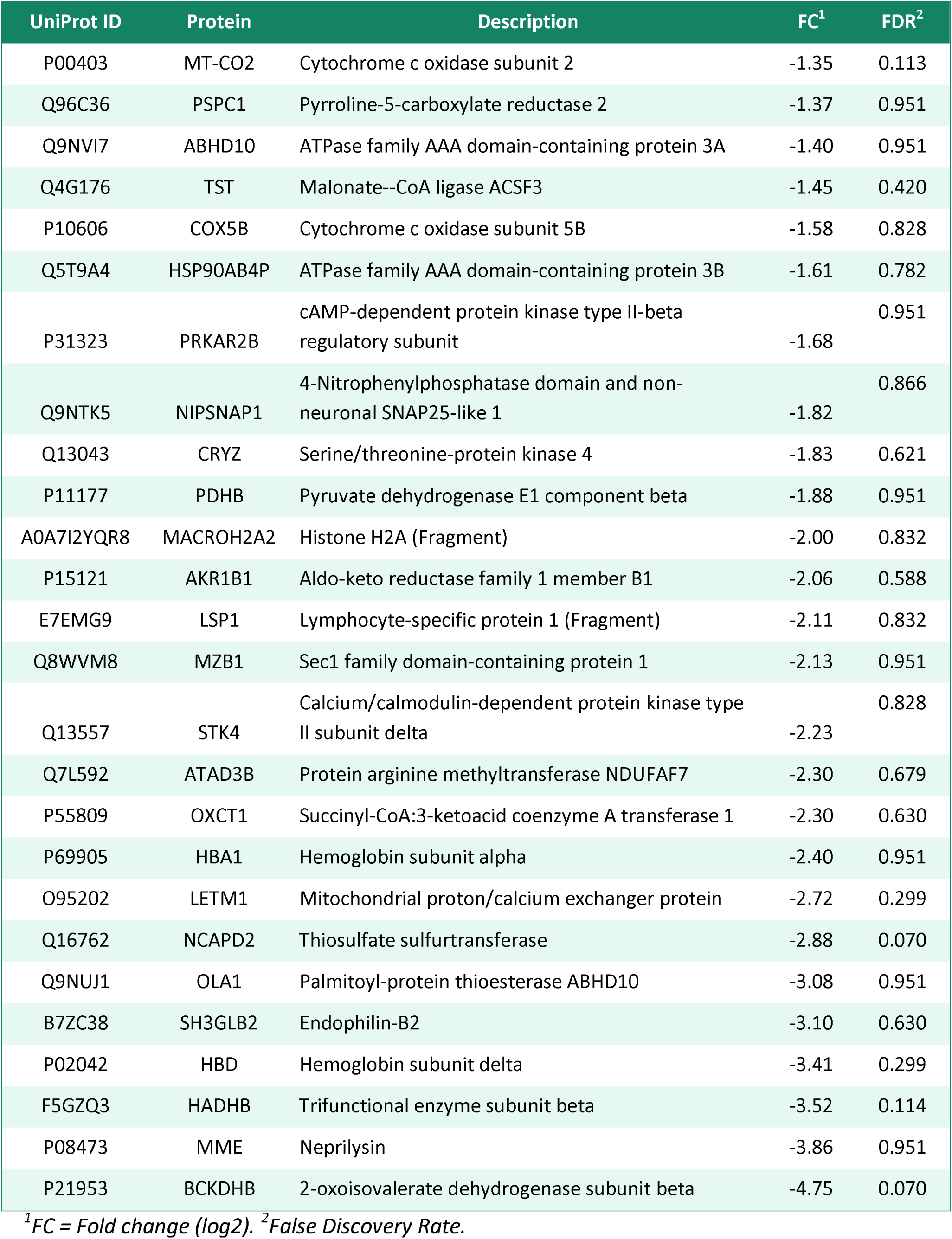
List of proteins significantly up- and down-regulated in ΔBART9_sg2 mutant compared to the Akata-EBV/Cas9 wild-type.

## 4. Discussion

In this study, we sought to investigate the effects of knocking down miRs BART 7 and 9 in lymphoid cells constitutively infected by EBV to get insights into the importance of these viral miRNAs in lymphomas associated with EBV infection. The influence and mechanisms of EBV-encoded miRs in associated lymphomas remain largely unknown. We have screened human lymphomas and lymphoblastoid cell lines by RT-qPCR and found significant heterogeneity in the expression of EBV miRs BART 7 and 9 (Supplementary Figure S2).These differences are presumably attributed to inherent distinctive biological features of these cell lines, as the expression levels of miRs BART vary across EBV-associated tumors, showing specific profiles due to growth stimuli and tumor microenvironment[6,21]. For instance, in a study comparing the molecular profile of endemic, sporadic, and immunodeficiency-associated BL, EBV-positive tumors that express high viral miRs levels were correlated with increased activation of proliferation pathways[22].

To study the effects of miRs BART7 and BART9 abrogation on Akata EBV-positive cells, we have created mutant cell lines by editing the EBV genome inside infected cells using CRISPR/Cas9. We were able to produce EBV mutants at a high genome editing efficiency, validated by T7EI mutation assays, Sanger sequencing, and decomposition analysis. Although CRISPR/Cas9 edition is commonly used to knockout genes, the complete abrogation of EBV miRs BART7 and BART9 in the current experimental setup could not be achieved. We observed a drastic reduction in miRs BART7 and BART9 expression in Akata-EBV/Cas9 mutants as assessed by RT-qPCR and luciferase reporter assays. Our results are similar to that found in the literature, where editing EBV multicopy episomes using CRISPR/Cas9 always yielded a low percentage of unedited episomes[23]. Nonetheless, our results highlight that using CRISPR/Cas9 to knock down the expression of EBV miRs BART is still a more robust and permanent option than other techniques currently employed to repress miRs expression, such as methods based on RNA interference technologies[24].

The Akata-EBV/Cas9 mutants with knockdown of EBV miRs BART7 and BART9 exhibited a significant reduction in cell viability and proliferation rates compared to the Akata WT controls. Our findings agree with the literature for NPC and BL-derived cell lines, where expression of BART7 and BART9 are associated with increased cell proliferation and resistance to apoptosis[25,9,26]. In addition, we also found that both BART7 and BART9 mutants display an increased expression of the viral genes encoding the lytic proteins ZTA and gp350 (BLZF1 and BLLF1, respectively). EBV BART9 was reported to attenuate and regulate the EBV lytic cycle via transcriptional inhibition of several components of the BCR and FOXO3 signaling pathway, showing striking sequence homology to the human miRNA miR-141 (hsa-miR-141)[27,28]. Our results support the hypothesis that the expression of BART7 and BART9 are associated with regulating cell proliferation, apoptosis and maintaining the viral latency in B-cells.

The global proteomic analysis performed in this study showed that the suppression of EBV BART7 significantly decreased the expression of ribosomal proteins, including proteins within the ubiquitin-proteasome system (UPS) such as RPS27A and UBA52, as well as the translational initiation factors RPS21 and EIF3L. Curiously, the expression of RPS27A was reported to induce cell proliferation mediated by LMP1, acting as a regulator of the viral oncoprotein by stabilizing it in a proteosome-dependent pathway^32^. Regulation of LMP1 by EBV miRs may have a vital role in increasing the aggressiveness of EBV-associated cancers by regulating signaling pathways associated with cell survival, cell proliferation, and tumor progression[30].

The ubiquitin/proteasome pathway is a complex proteolytic system regulating several critical cellular processes, including signal transduction, cell cycle progression, apoptosis, and MHC class I loading complex.[31] EBV can exploit this pathway by either inhibiting proteasomal degradation to avoid the loss of critical viral proteins or inducing the degradation of unwanted cellular proteins ^32,33,34^. Because the ubiquitin/proteasome pathway is pivotal for generating antigenic peptides of viral proteins for presentation to T cytotoxic cells mediated by MHC-I molecules, its inhibition by EBV (mediated by EBNA-1, for instance) favors immune evasion of EBV-infected cells[35].

Our study also shows that the suppression of BART7 significantly increased the expression of RBPs, including proteins directly associated with ribosome biogenesis, such as STRAP, WDR12, GAR1, and ISG20. The pathway enrichment analysis showed several hits for ribonucleoprotein complex binding, RNA binding, and nucleic acid binding. RBPs are master regulators of mRNA biogenesis and maturation, exerting crucial activities in normal hematopoiesis and hematological malignancies[37–39]. During the lytic cycle, EBV recruits several host’s RBPs and ribosomal proteins, coopted for the translation, maturation, and regulation of viral proteins.[40,41] In our model, suppression of BART7 led to a significant increase in the expression of the Zta and gp350 lytic genes. Thus, it is plausible that BART7 may selectively target human RBPs to tune in the lytic replication to maintain persistent infection.

Finally, knocking down BART9 significantly reduced the expression of several mitochondrial enzymes, including ACSF3, ALDH5A1, HADHB, IDH2, OXCT1, and PDHB, particularly enriched in fatty acid metabolism, mediated by oxidoreductase activity pathways. Reprogramming of fatty acid metabolism occurs in hematological malignancies and favors tumorigenesis and cell survival. In this regard, knockdown of HADHB was reported to suppress the proliferation of LCL-K and MD901 cells; also, IDH2 and OXCT1 overexpression was correlated to increase cell survival, proliferation, autophagy, and chemoresistance in different cancer cell models[42–45]. Our findings disclose a possible role of EBV BART7 in regulating lipid metabolism and support the hypothesis that EBV products may subvert these metabolic pathways to promote B-cell survival and proliferation[46,47].

In conclusion, our findings suggest potential roles for EBV miRs BART7 and BART9 in Akata EBV-positive cells, increasing cell proliferation and viability, modulating the lytic cycle and the expression of immune checkpoints, and hijacking several critical cellular pathways, including RBPs, ubiquitin/proteasome and fatty acid metabolism.

## Acknowledgments

This study was sponsored by the Brazilian public agencies CNPq, CAPES, and FAPESP, which awarded scholarships to BFRC (CNPQ DR 142294/2017-9; FAPESP DR 2017/20352-0, FAPESP BEPE DR 2019/03804-0), VLR (CAPES 88887.466765/2019-00) and research funds to DEO (FAPESP Grant 2017/23393-9). We thank Professors Benjamin E. Gewurz and Ethel Cesarman for kindly providing cells and reagents used in this study.

## Data availability statement

The data that support the findings of this study are openly available in OSF at https://osf.io/hksuf. The mass spectrometry data have been uploaded to the MassIVE Repository (Computer Science and Engineering, University of California, San Diego, USA) and is available at ftp://massive.ucsd.edu/MSV000090486 with the dataset identifier MSV000090486.

