## Supplementary Material for "Epstein-Barr virus miR-BARTs 7 and 9 modulate viral cycle, cell proliferation and proteomic profiles in Burkitt lymphoma"

**Table S1 –** STR profiles of the cell lines used in the study

| Cell Line  (RRID) | Description | STR profile |
| --- | --- | --- |
| Akata-Cas9^a^  (CVCL_1856) | Burkitt lymphoma cells, stably infected by EBV (Latency type I) and expressing of Cas9. | AMEL: X,X; CSF1PO: 11; D5S818: 11,12; D7S820: 10,12; D13S317: 8,12; D16S539: D21S11: 30,32.2, 10,11; TH01: 9; TPOX: 8,11; vWA: 14,16 |
| BC-1^b^  CVCL_1079 | Primary effusion lymphoma from ascitic fluid; stably infected by EBV (Lat I); coinfected by KSHV and HIV. | AMEL: X, Y; CSF1PO: 10,11; D13S317: 8; D16S539: 12, 13; D21S11: 28, 30; D5S818: 11, 12; D7S820: 11; TH01: 9.3; TPOX: 8, 9; vWA: 16,20 |
| BC-2^b^  CVCL_1856 | Primary effusion lymphoma from pleural fluid; stably infected by EBV (Lat I); coinfected by KSHV and HIV. | AMEL: X,Y; CSF1PO: 10,12; D13S317: 11,12; D16S539: 11,12; D21S11: 29,30; D5S818: 11,13; D7S820: 8,9; TH01: 7; TPOX: 10,11; vWA: 14,19 |
| BC-3^b^  CVCL_1080 | Primary effusion lymphoma from pleural fluid; EBV negative, infected by KSHV and HIV. | AMEL: X CSF1PO: 11, 12; D13S317: 11; D16S539: 12; D21S11: 29, 30; D5S818: 11,12; D7S820: 10, 12; TH01: 6, 9; TPOX: 8, 11; vWA: 14,18 |
| IBL-1^b^  CVCL_9638 | Diffuse large B cell lymphoma; stably infected by EBV (Lat II/III); coinfected by HIV. | AMEL: X,Y; CSF1PO: 12,12; D5S818: 11,12; D7S820: 12,13; D13S317: 11,11; D16S539: 9,12; D21S11: 28,29; TH01: 9,9.3; TPOX: 8,11; vWA: 17,19 |
| Jiyoye^b^  CVCL_1317 | Burkitt lymphoma; stably infected by EBV (Lat I). | AMEL: X,Y; CSF1PO: 10,11; D5S818: 12,12; D7S820: 8,10; D13S317: 12,12; D16S539: 10,11; D21S11: 28,36; TH01: 7,9; TPOX: 6,8; vWA: 15,19 |
| MutuI-Cas9^a^  CVCL_7202 | Burkitt lymphoma, stable expression of Cas9 and infection by EBV (Lat I). | AMEL: x,y; CSF1PO: 10; D5S818: 12,14; D7S820: 8,10; D13S317: 12,13; D16S539: 13; D21S11: 30,32.2; TH01: 7; TPOX: 8,11; vWA: 15,17 |
| P3HR1-Cas9^a^  CVCL_2676 | Burkitt lymphoma, stable expression of Cas9 and infection by EBV (Lat I) | AMEL: X,Y; CSF1PO: 10,11; D5S818: 12,12; D7S820: 8,10; D13S317: 12,12; D16S539: 10,11; D21S11: 28,36; TH01: 7,9; TPOX: 6,8; vWA: 15,19 |

^1^EBV latency type (I, II or III); ^a^A gift from Benjamin Gewurz at the Harvard Medical School; ^b^A gift from Ethel Cesarman at the Weill Medical College, Cornell University.

**Table S2 –** Reagents and resources

| Reagent | Source | Cat. No. |
| --- | --- | --- |
| 2-Iodoacetamide | Sigma-Aldrich | 161125 |
| 3500 Series Genetic Analyzer | Thermo Fisher Scientific | 4406017 |
| Ammonium bicarbonate | Sigma-Aldrich | A6141 |
| Ampicillin sodium salt | Sigma-Aldrich | A0166 |
| Bio-Rad Protein Assay Kit | Bio-Rad | 5000001 |
| Blasticidin S HCl (10 mg/mL) | Thermo Fisher Scientific | A1113903 |
| BsmBI-v2 Restriction Enzyme | New England Biolabs | R0739S |
| CellTiter 96® AQueous One Solution | Promega | G3582 |
| cOmplete™, EDTA-free Protease Inhibitor Cocktail | Roche | 11873580001 |
| Countess™ 3 FL Automated Cell Counter | Thermo Fisher Scientific | AMQAF2000 |
| Dithiothreitol | Bio-Rad | 161-0611 |
| DMEM, powder, high glucose | Thermo Fisher Scientific | 12100061 |
| DNase I, Amplification Grade | Sigma-Aldrich | AMPD1-1KT |
| Dual-Glo® Luciferase Assay System | Promega | E2920 |
| Fetal Bovine Serum | Sigma-Aldrich | F7524 |
| Filtropur S, PES, pore size: 0.45 µm | Sarstedt Inc | 83.1826.001 |
| Gel Loading Dye, Orange (6X) | New England Biolabs | B7022S |
| GenePrint® 10 System | Promega | B9510 |
| Geneticin™ Selective Antibiotic (G418 Sulfate) | Thermo Fisher Scientific | 10131035 |
| Gentamicin (10 mg/mL) | Thermo Fisher Scientific | 15710064 |
| GloMax® Discover Microplate Reader | Promega | GM3000 |
| GoTaq® qPCR Mix | Promega | A6001 |
| High-Capacity cDNA Reverse Transcription Kit | Thermo Fisher Scientific | 4374966 |
| LB Agar, powder (Lennox L agar) | Thermo Fisher Scientific | 22700041 |
| Lipofectamine 3000 transfection reagent | Thermo Fisher Scientific | L3000015 |
| Luria Broth Base (Miller's LB Broth Base)™, powder | Thermo Fisher Scientific | 12795084 |
| One-Shot Stbl3 Chemically Competent E. coli | Thermo Fisher Scientific | C737303 |
| Opti-MEM™ I Reduced Serum Medium | Thermo Fisher Scientific | 31985070 |
| pGL4.73[hRluc/SV40] Vector | Promega | E6911 |
| Pierce™ C18 Spin Columns | Thermo Fisher Scientific | 89870 |
| Platinum™ Taq DNA Polymerase High Fidelity | Thermo Fisher Scientific | 11304029 |
| Puromycin Dihydrochloride | Thermo Fisher Scientific | A1113803 |
| QIAquick PCR & Gel Cleanup Kit | Qiagen | 28506 |
| Quick Start Bovine Serum Albumin Standard Set | Bio-Rad | 5000207 |
| RapiGest SF | Waters | 186001861 |
| Reverse Mass DNA Ladder | New England Biolabs | N3240S |
| RNeasy Mini Kit | Qiagen | 74104 |
| RPMI 1640 Medium, powder | Thermo Fisher Scientific | 31800105 |
| Sequencing Grade Modified Trypsin | Promega | V5111 |
| T4 DNA Ligase | New England Biolabs | M0202S |
| T4 DNA Ligase Reaction Buffer | New England Biolabs | B0202S |
| T7 Endonuclease I | New England Biolabs | M0302S |
| TaKaRa PCR Mycoplasma Detection Set | Takara Bio Inc | 6601 |
| Trifluoroacetic acid | Sigma-Aldrich | 302031 |
| TRIzol™ Reagent | Thermo Fisher Scientific | 15596026 |
| Vacufuge® Concentrator Plus | Eppendorf | EP5305000169 |
| Vibra-Cell™ VCX 750 | Sonics & Materials Inc. | VCX 750 |
| Wizard® Genomic DNA Purification Kit | Promega | A1120 |

**Table S3 –** Sequences of oligonucleotides used in the study

| Name | Sequences (5’-3’) |
| --- | --- |
| sgRNA sequence cloned into LentiGuide-Puro | |
| BART7_sg1 | F: CACCGTTGTTTCATAGTCAAGGTCC / R: AAACGGACCTTGACTATGAAACAAC |
| BART7_sg2 | F: CACCGCATAGTCAAGGTCCAGGATC/ R: AAACGATCCTGGACCTTGACTATGC |
| BART9_sg1 | F: CACCGCTGAATTGGAAACAGTAACT / R: AAACAGTTACTGTTTCCAATTCAGC |
| BART9_sg2 | F: CACCGAAGTTACTGTTTCCAATTCA / R: AAACTGAATTGGAAACAGTAACTTC |
| Sequencing of LentiGuide-Puro constructs | |
| hU6.F | GAGGGCCTATTTCCCATGATT |
| Luciferase Reporter Assays | |
| psiCheck-2-BART7.3p | AAACCATCATAGTCCAGTGTCCAGGGTCTAGAC  TCGAGTCTAGACCCTGGACACTGGACTATGATGGTTT |
| psiCheck2 BART7.3p SCR | AAACGTAAGACTAGGCCATCGTGCTCTCTAGAC  TCGAGTCTAGAGAATACTCCGTATCGTGAGAAGCGTTT |
| psiCheck-2-BART9.3p | AAACTAACACTTCATGGGTCCCGTAGTTCTAGAC  TCGAGTCTAGAACTACGGGACCCATGAAGTGTTAGTTT |
| psiCheck-2-BART7.3p SCR | AAACGCTTCTCACGATACGGAGTATTCTCTAGAC  TCGAGTCTAGAGAATACTCCGTATCGTGAGAAGCGTTT |
| Sequencing of EBV miRs BARTs 7 and 9 within the EBV genome | |
| BART7_seq | F: ATTCTGTTCTATGACCCCGT / R: CTAGGAAACCGTAATCAGTG |
| BART9_seq | F: GTATTTTCCCATCAGCACCT / R: ACCATGACTTTGTAACCGAG |
| EBV lytic cycle evaluation by RT-qPCR | |
| Zta (BZLF1) | F: TACAAGAATCGGGTGGCTTC / R: GCACATCTGCTTCAACAGGA |
| gp350 (BLLF1) | F: TGTTACAGTGACTGCCTTTTGGG / R: GGTGTCCCCGAGGTGAGAGT |
| Analysis of Immune-checkpoint molecules by RT-qPCR | |
| Tim3 | F: CTTTCCAAGGATGCTTACCAC / R: CAGATCCCTGCTCCGATGTA |
| PDL1 | F: GCCCCATACAACAAAATCAACC / R: GCTTGTCCAGATGACTTCGG |

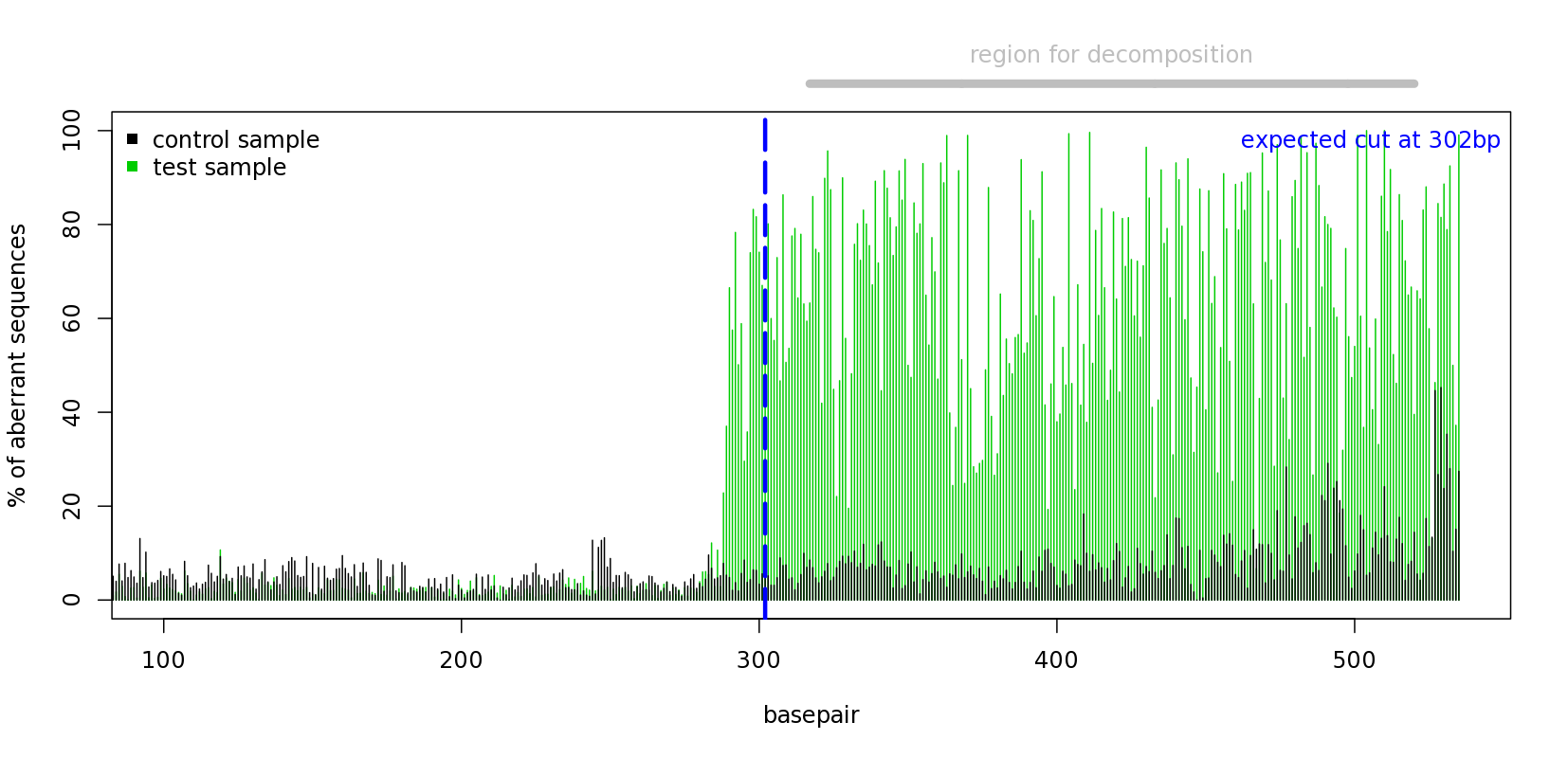

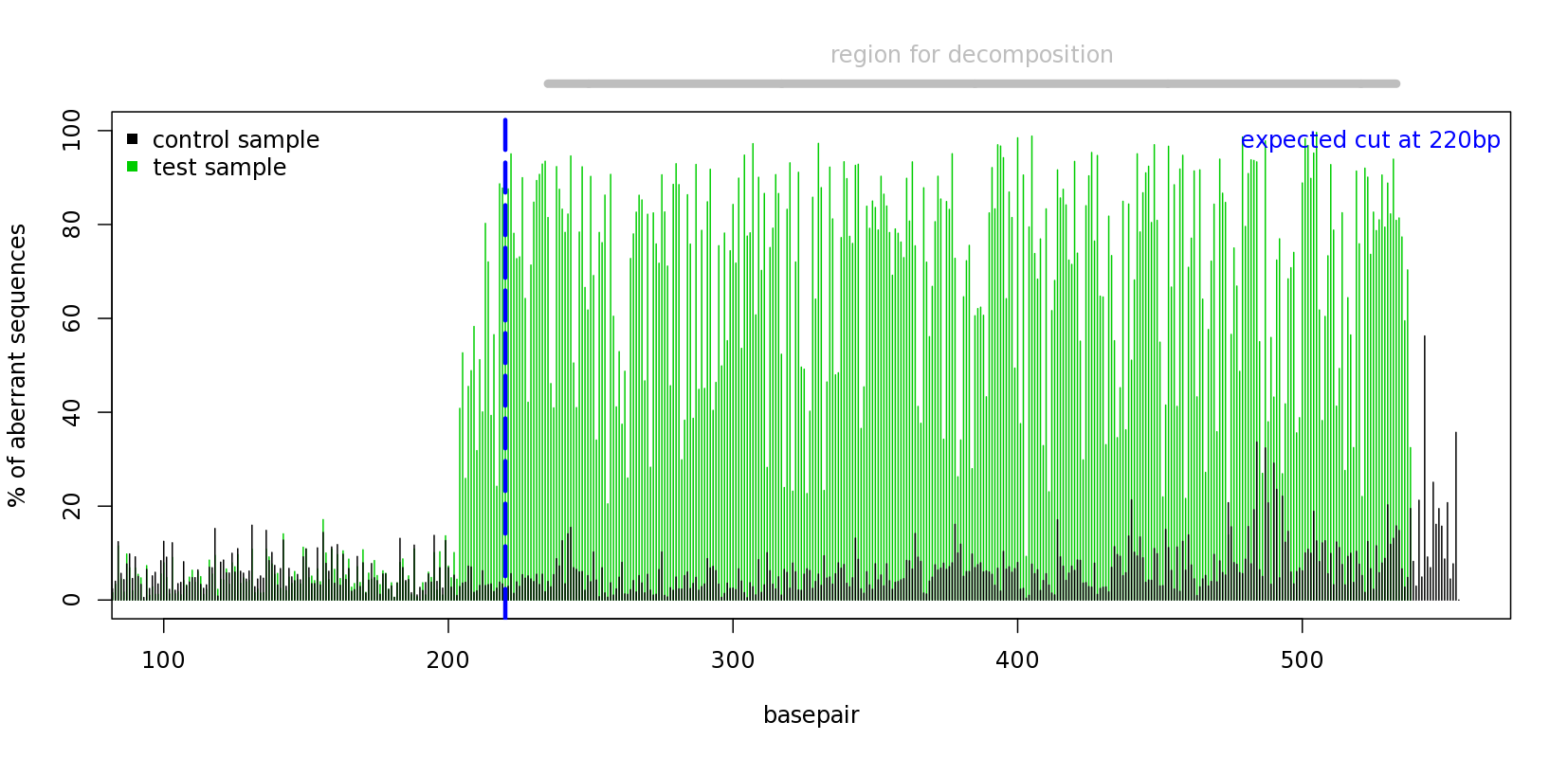

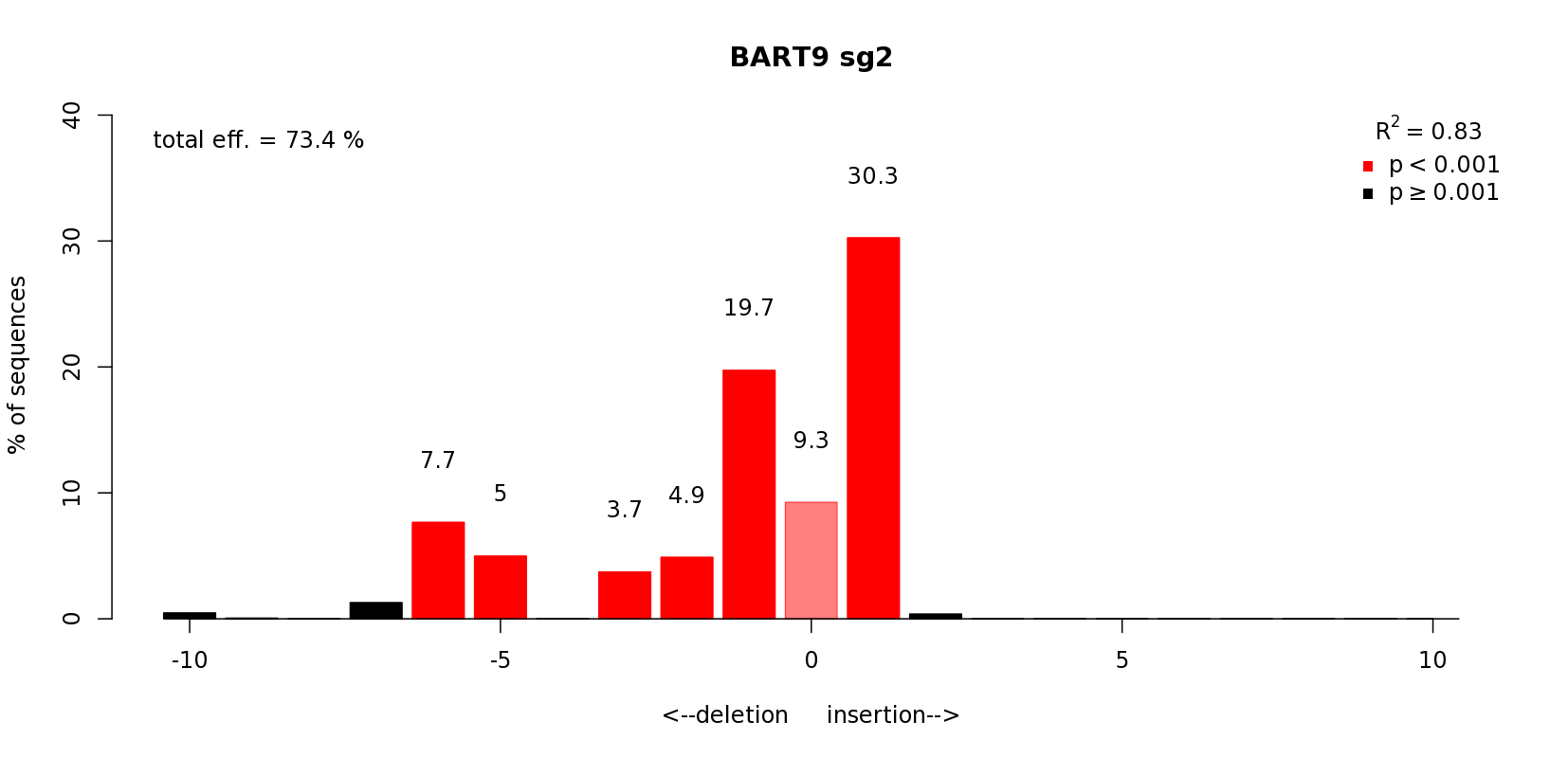

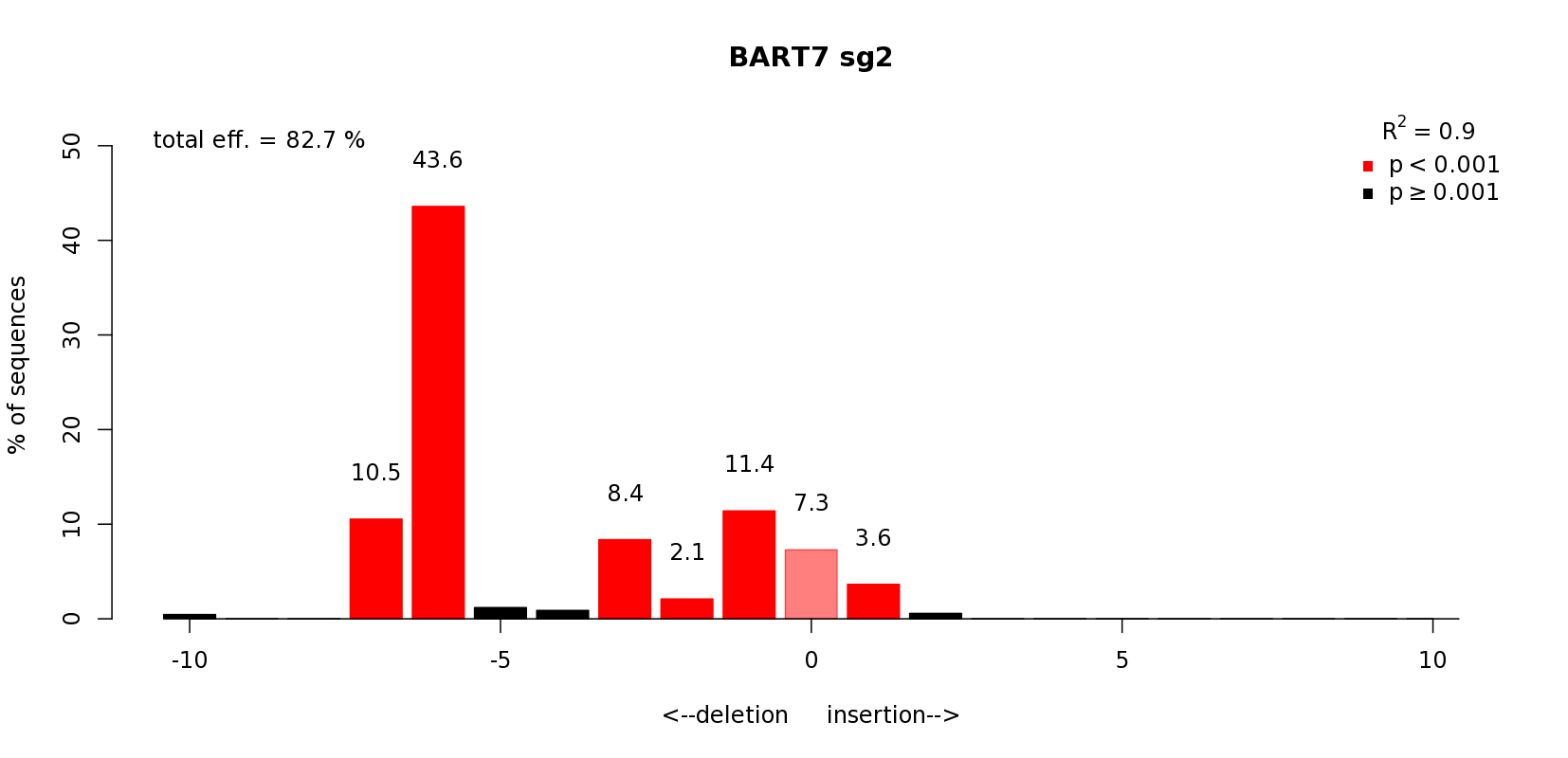

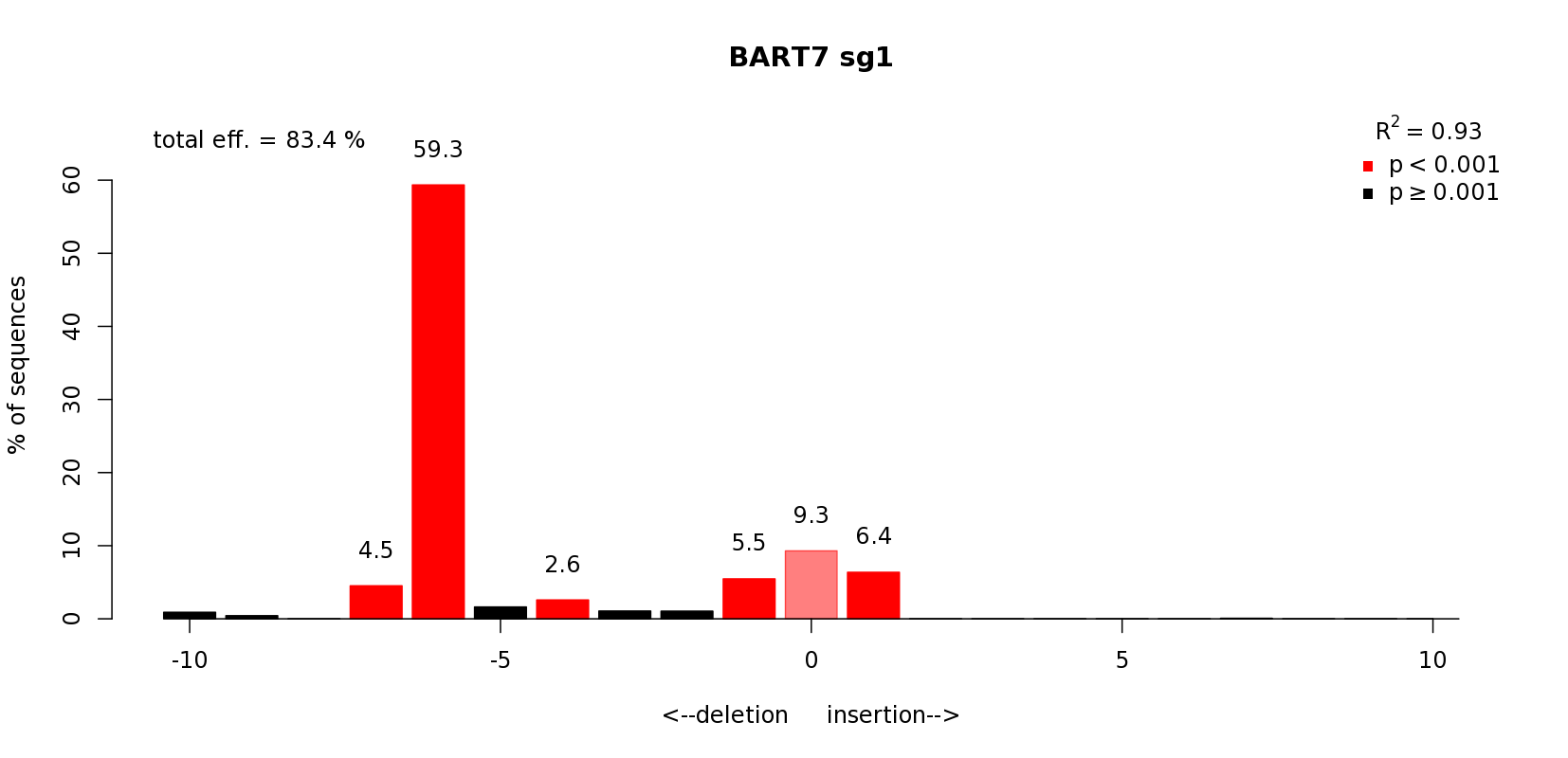

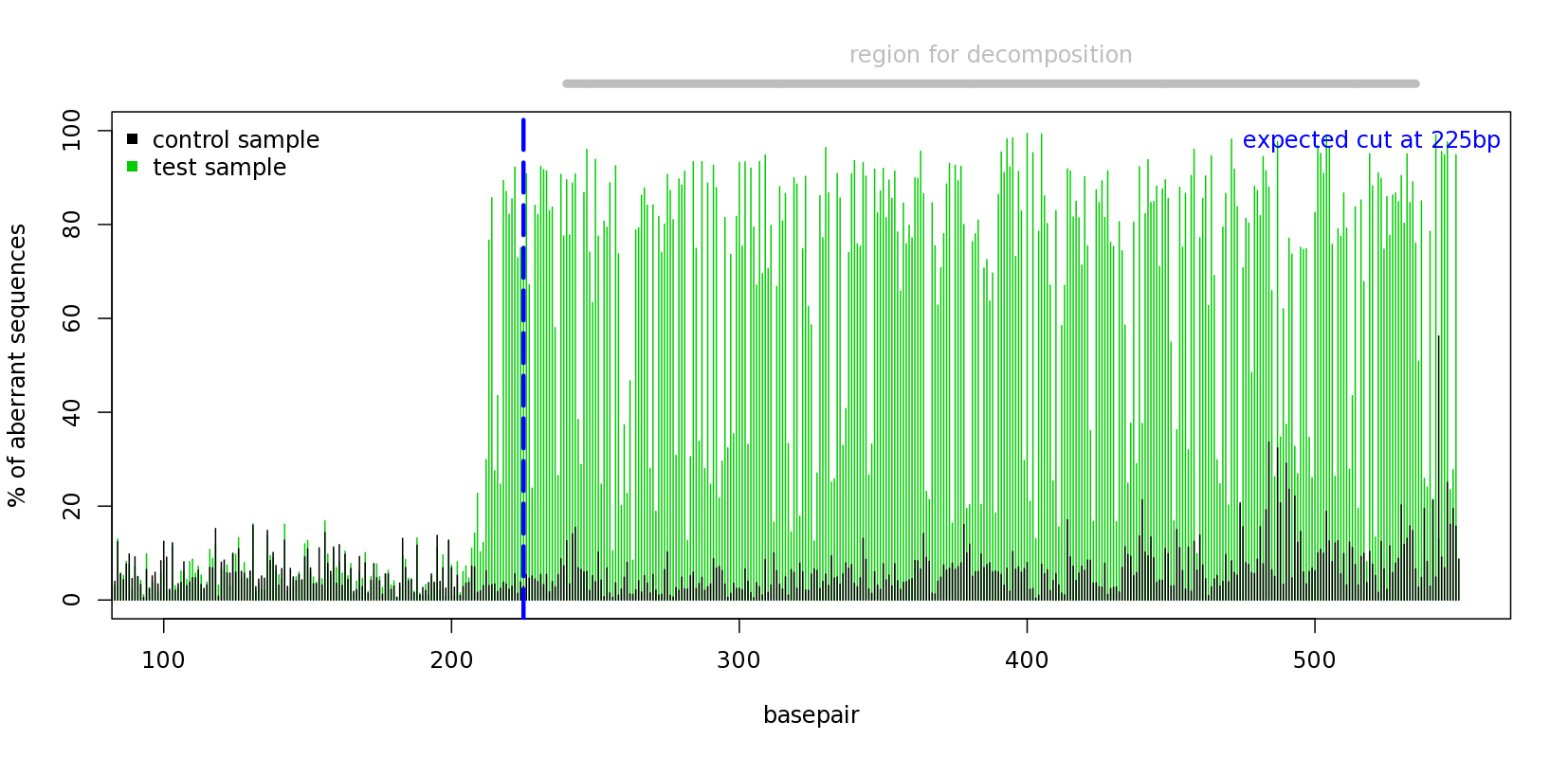

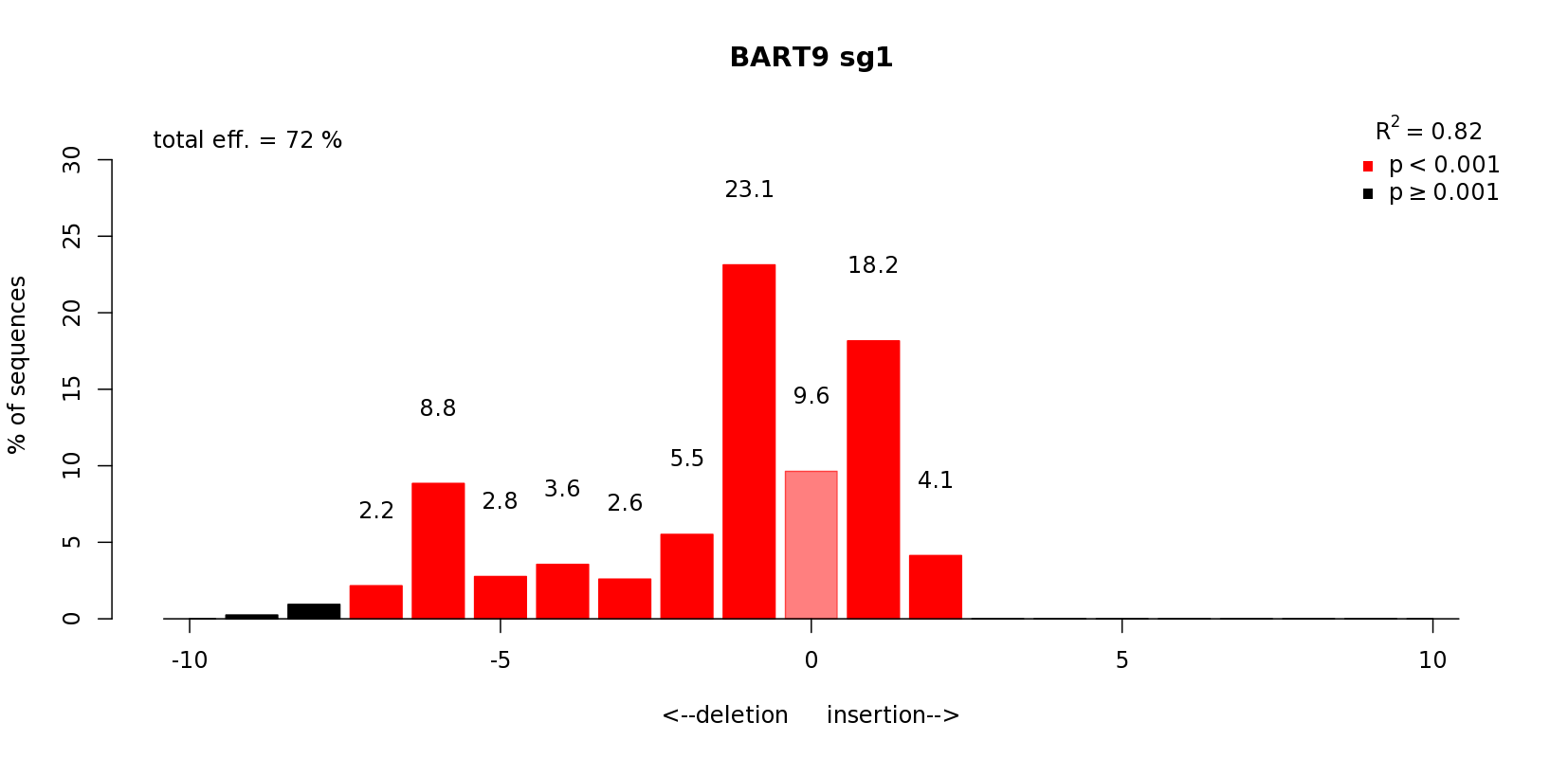

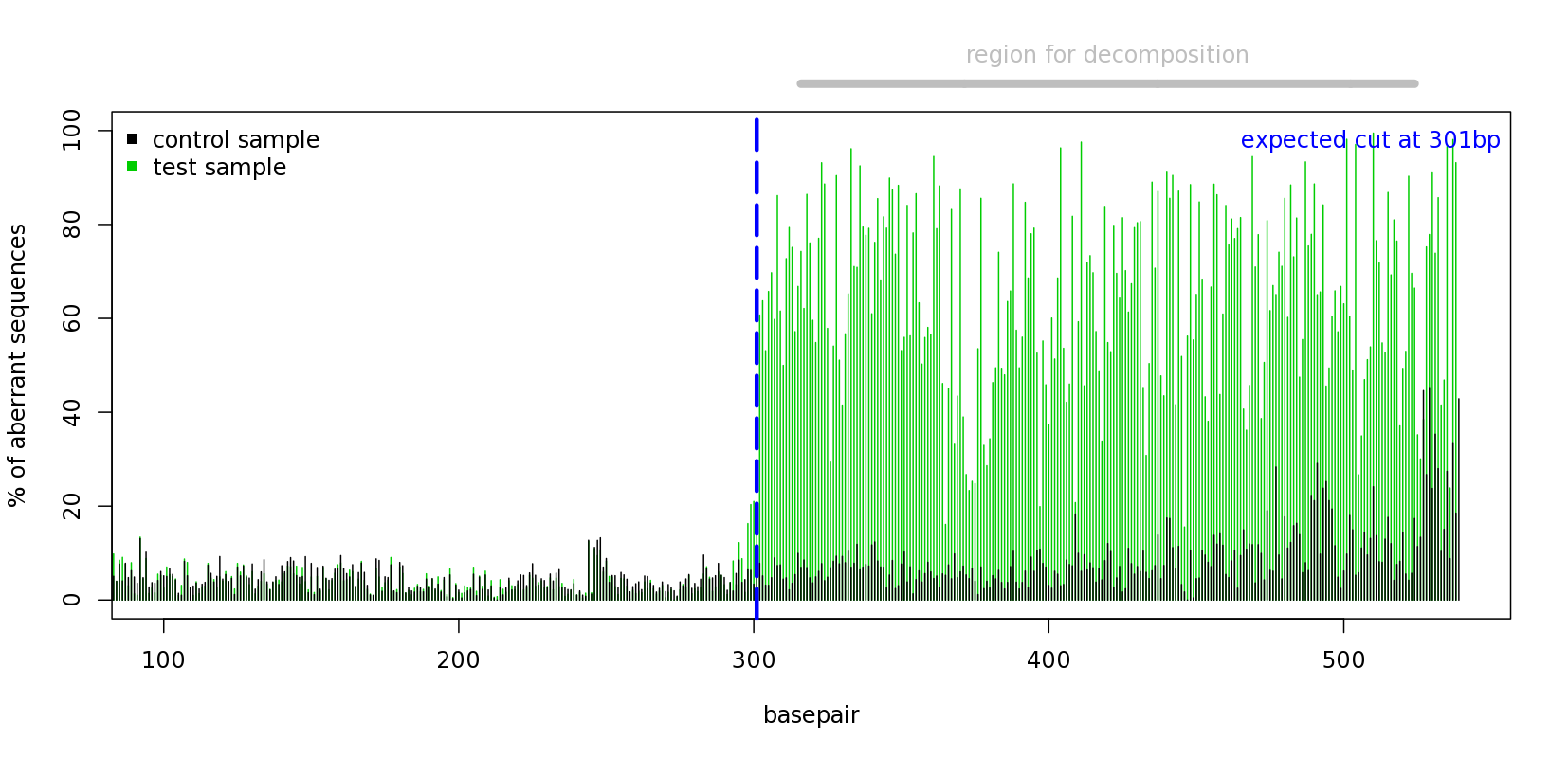

**A**

**B**

**C**

**D**

**Supplementary Figure S1 –** Genome editing efficiency and INDEL profiles of Akata mutants using the Tracking of Indels by Decomposition (TIDE) analysis. Sequences of (**A**) ΔBART7_sg1, (**B**) ΔBART7_sg2, (**C**) ΔBART9_sg1 and (**D**) ΔBART9_sg2, compared to Akata WT.

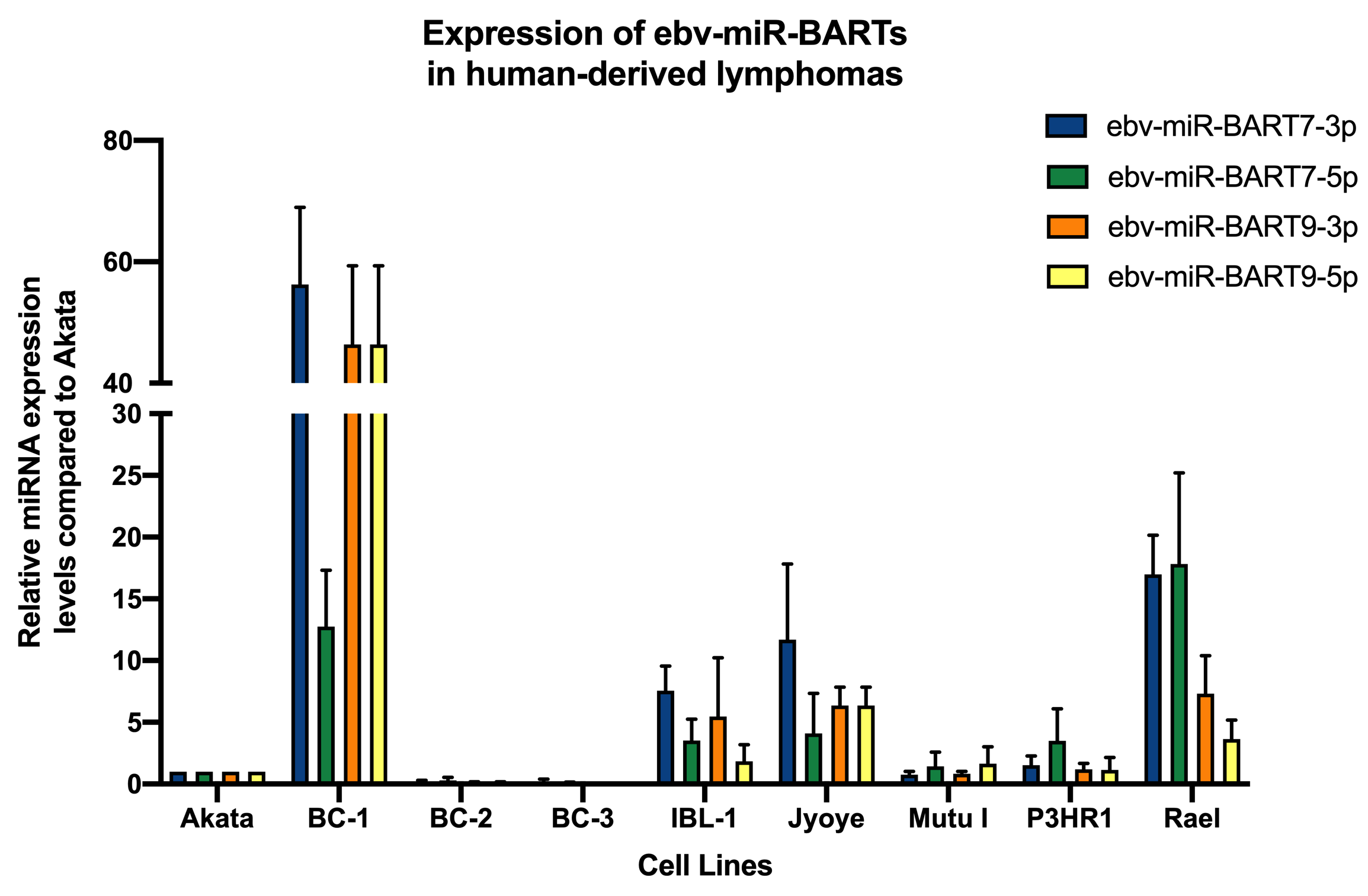

**Supplementary Figure S2–** Expression of EBV miRs BARTs 7 and 9 in human-derived lymphomas associated with EBV infection. BC3 (EBV-negative PEL) and Akata EBV/Cas9 are negative control and reference (expression = 1), respectively. The human miRNAs Snord47 and Snord48 were used as endogenous controls. Data expressed as mean ± SD.
